# Quantitative Mapping of Human Hair Graying and Reversal in Relation to Life Stress

**DOI:** 10.1101/2020.05.18.101964

**Authors:** Ayelet Rosenberg, Shannon Rausser, Junting Ren, Eugene Mosharov, Gabriel Sturm, R Todd Ogden, Purvi Patel, Rajesh Kumar Soni, Clay Lacefield, Desmond J Tobin, Ralf Paus, Martin Picard

## Abstract

Hair graying is a universal hallmark of aging believed to be linked to psychological stress. Here we develop a novel approach to quantitatively profile natural graying events along individual human hair shafts, resulting in a quantifiable physical timescale of hair pigmentation patterns (HPPs). Using this approach, we quantify rare events of white/gray hairs that naturally regain pigmentation within days to weeks, thereby quantitatively defining the reversibility of graying in healthy, unmedicated individuals. Proteomic analysis shows that graying is marked by the upregulation of proteins related to energy metabolism, mitochondria, and antioxidant defenses. Combining hair pigmentation profiling and proteomics at the single hair level, we also report hair graying and its reversal occurring in parallel with behavioral and psychological stressors. A computational simulation suggests a threshold-based mechanism for the temporary reversibility of graying. Quantitatively mapping HPPs in humans provides an opportunity to longitudinally examine the influence of life exposures on biological aging.

## Main Text

Hair graying is a ubiquitous, visible, and early feature of human biological aging (*1*). The time of onset of hair graying varies between individuals, as well as between individual hair follicles, based on genetic and other biobehavioral factors (*2, 3*). But most people experience depigmentation of a progressively large number of hair shafts (HSs) from their third decade onward, known as achromotrichia or canities (*4*). The color in pigmented HSs is provided by melanin granules, a mature form of melanosomes continuously supplied to the trichocytes of the growing hair shaft by melanocytes of the hair follicle pigmentary unit (HFPU) (*1*). Age-related graying is thought to involve melanocyte stem cell (MSC) exhaustion (*5*), neuroendocrine alterations (*6*), and other factors, with oxidative damage to the HFPU likely being the dominant, initial driver (*6–8*). While loss of pigmentation is the most visible change among graying hairs, depigmented hairs also differ in other ways from their pigmented counterparts (*9*), including in their growth rates (*10*), HF cycle, and other biophysical properties (*11*). Hair growth is an energetically demanding process (*12*) relying on aerobic metabolism in the HF (*13*). Melanosome maturation also involves the central organelle of energy metabolism, mitochondria (*14, 15*). Moreover, mitochondria likely contribute to oxidative stress within the HF (*16*), providing converging evidence that white hairs may exhibit specific alterations in mitochondrial energy metabolism.

Although hair graying is generally considered a progressive and irreversible age-related process, with the exclusion of alopecia areata (*17*), various cases of drug- and mineral deficiency-induced depigmentation or repigmentation of hair have been reported (*18–24*) reflecting the influence of environmental inputs into HFPU function (*25*). While spontaneous repigmentation can be pharmacologically-induced, its natural occurrence in unmedicated individuals is rare and has only been reported in a few single-patient case studies (*26–30*). A proposed mechanism for such repigmentation events involve the activation and differentiation of a subpopulation of immature melanocytes located in a reservoir outside of the hair follicle bulb in the upper outer root sheath (*11*). However, the reversal of hair graying has not been quantitatively examined in a cohort of healthy adults, in parallel with molecular factors and psychosocial exposures.

The influence of psychological stress on hair pigmentation is a debated but poorly documented aspect of hair graying. In humans, psychological stress accelerates biological aging as measured by telomere length (*31, 32*). In mice, psychological stress and the stress mediator norepinephrine acutely causes depigmentation (*33*), but in mice graying is an irreversible phenomenon driven by a depletion of melanocyte stem cells (*8*). In humans, where HF biology differs significantly from rodents (*30*), stress-induced graying and its reversal remain insufficiently understood. The paucity of quantitative data in humans is mostly due to the lack of sensitive methods to precisely correlate stressful psychobiological processes with hair pigmentation and graying events at the single-follicle level.

Here we describe a digitization approach to map hair pigmentation patterns (HPPs) in single hairs undergoing graying and reversal transitions, examine proteomic features of depigmented white hairs, and illustrate the utility of the HPP approach to interrogate the association of life stress and hair graying in humans. Because previous literature suggests that rare pigmentation events are more likely to occur in the early stages of canities (*11*), the current study focuses primarily on pigmentation events in young to middle-aged participants. Finally, we develop a computational model of hair graying to explore the potential mechanistic basis for stress-induced graying and reversibility on the human scalp hair population, which could potentially serve as a resource for the *in silico* modelling of macroscopic aging events in human organs/tissues.

## Results

### Mapping hair pigmentation patterns (HPPs)

To overcome the lack of methodology to map pigmentary states and age-related graying transitions, we developed an approach to digitize HPPs at high resolution across the length of single human HSs. Combined with known hair growth rates on the scalp (~1.0-1.3 cm per month (*34*)), this approach provides a quantifiable, personalized live bioarchive with the necessary spatio-temporal resolution to map individualized HPPs and graying events along single hairs, and to link HPPs to specific moments in time with unprecedented accuracy. Using this methodology, similar to dendrochronology where tree rings represent elapsed years (*35*), hair length reflects time, and the HS length is viewed as a physical time scale whose proximal region has been most recently produced by the HF, and where the distal hair tip represents weeks to years in the past, depending on the HS length.

To examine HPPs in human hairs, we plucked, imaged, digitized, and analyzed hairs (n=397) from 14 healthy donors (**Figure 1A**) (see Methods for details). Three main pigmentation patterns initially emerged from this analysis: i) Hairs with constant high optical density (*Dark*), ii) Hairs with constant low optical density (*White*); iii) Initially dark hairs that undergo a sharp graying transition from dark to white over the course of a single growing anagen phase of the hair follicle growth cycle (*Transition*) (**Figure 1B-C**). Dark-to-white transitions demonstrate the existence of rapid depigmentation events within a single anagen hair cycle (*36, 37*). We confirmed that compared to dark hairs still harboring their “young” pigmentary state, the HFPU of “aged” white HFs from either African American or Caucasian individuals are practically devoid of pigment (**Figure 1D**), which is consistent with the finding of previous studies (*38*). Whereas dark hairs contain melanin granules dispersed throughout the hair cortex when observed by electron microscopy, white hairs from the same individuals show a near complete (>98%) absence of melanin, with the few retained melanin granules, when present, being smaller, less dense, and at times vacuolated, a potential response to oxidative stress (*39*) (**Figure 1E-I**, see **Supplemental Figure S1** for high-resolution images of mature melanosomes). The digitization of HPPs thus reflects the presence of melanosomes within the HS, and rapid graying events are marked by the loss of melanosomes.

**Fig. 1.**
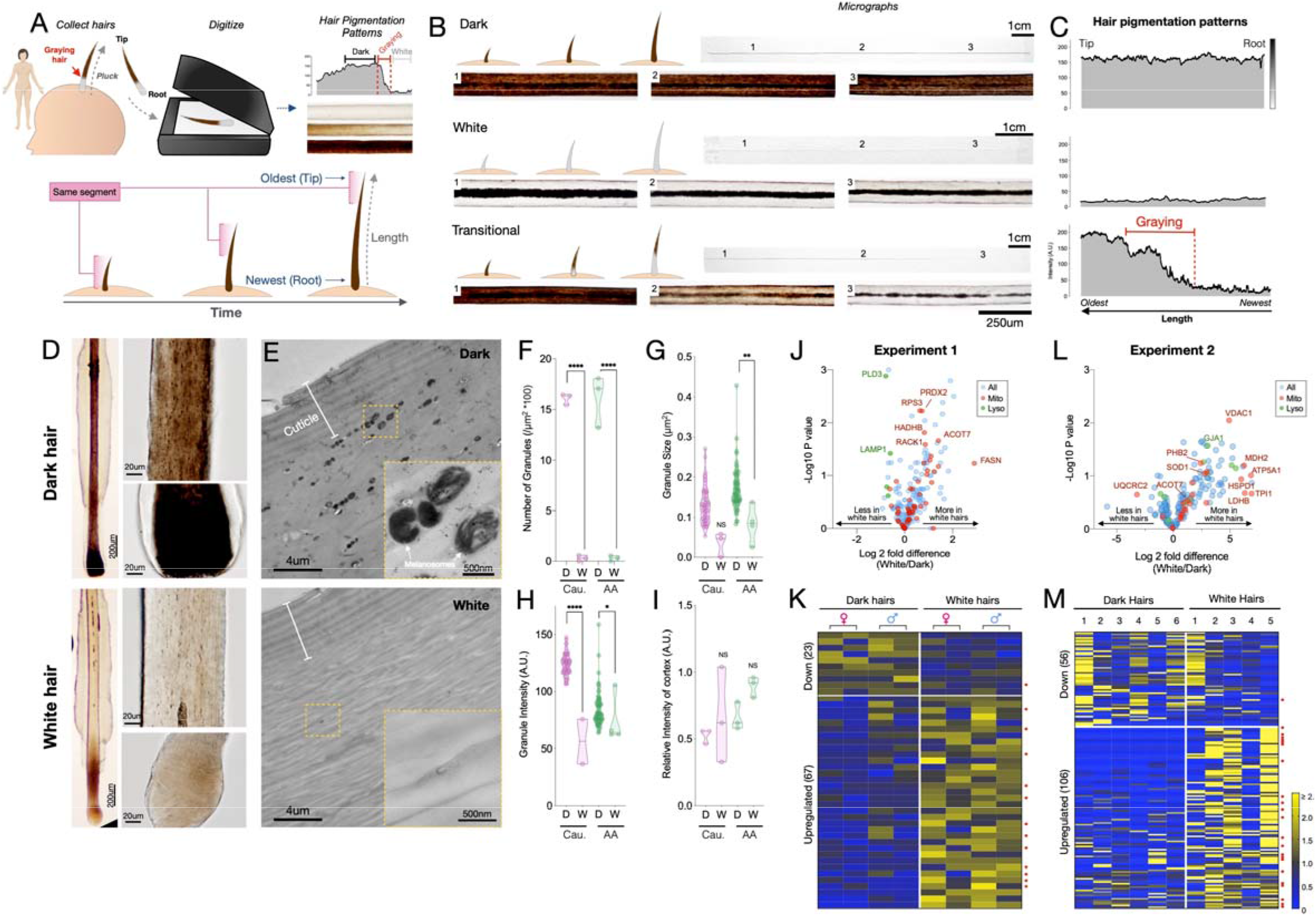
Quantitative analysis of human hair pigmentation patterns, graying, and associated proteomic changes. (**A**) Diagram illustrating hair growth over time, method of hair collection, digitization, and hair pigmentation pattern (HPP) methodology. (**B**) Dark, white, and hairs undergoing natural age-related transitions from the younger dark state to the older white state at macroscopic and microscopic resolution. (**C**) Digitized HPPs for the hairs shown in (B). (**D**) Bright field microscopy images of hair follicles from plucked dark (top-panel) and white hair (bottom-panel) from the same Caucasian male individual illustrating the loss of pigmentation in the hair follicle pigmentary unit (HFPU). (**E**) Electron microscopic images of dark (*top*) and white (*bottom*) scalp hairs from a Caucasian male showing absent melanin granules in white hairs. (**F**) Quantification from dark (D) and white (W) hairs (n=3 segments from each) from a Caucasian (Cau.) male and African-American (AA) male of melanin granule abundance, (**G**) size and (**H**) darkness. (**I**) Overall electron density of the hair matrix (excluding granules) (N.S.). **J**) Volcano plot comparing dark and white hair proteomes and (**K**) heatmap of down-(<0.8-fold) and up-regulated (>1.5-fold) proteins that were detected in all samples (n=90) for experiment 1 (duplicates of dark/white hairs from 1 female, 1 male, n=8 samples). (**L**) Volcano plot and (**M**) heatmap for all proteins detected in ≥3 samples (n=192) from experiment 2 (dark and white hairs from 6 individuals, n=6 dark and 5 white hairs). Proteins annotated as mitochondrial (Mitocarta2.0) and lysosomal (The Human Lysosome Gene Database) are highlighted. Red dots to the right of heatmaps indicate mitochondrial proteins. *P<0.05, **P<0.01, ****P<0.0001 from one-way ANOVA, Tukey’s multiple comparisons.

### Proteomic alterations in white hairs

To gain molecular insight into the graying process, we performed a comprehensive proteomic analysis comparing dark and white HS. Recent work suggests that depigmentation is associated with the upregulation of lipid synthesis enzymes in the HS (*40*). Moreover, in depigmented hairs, the abnormal diameter/caliber of the hair fiber, growth rate, presence/absence of HS medulla as well as the (dis)continuity and diameter of the medulla along the hair length (*41*) imply multiple potential proteomic alterations associated with depigmentation. In addition, melanogenesis involves high levels of reactive oxygen species, but dark HFs are equipped with multiple antioxidant mechanisms (e.g.,(*39*)). Thus, the proteomic features of HSs may provide interpretable information about molecular changes associated HF graying.

Protein extraction and LC-MS/MS methods optimized from a previous protocol (*42*) were used to process the unusually resistant proteinaceous matrix of the hair shaft and to handle the overly abundant keratin proteins over other potential proteins of interest (see Methods for details). Two independent experiments were performed. *Experiment 1*: matched dark and white hairs collected at the same time from two closely age- and diet-matched individuals (1 female and 1 male, both 35 years old, each dark and white HS measured twice, total n=8); and *Experiment 2 (validation):* n=17 hair segments from 7 different individuals (4 females and 3 males).

In the first experiment, we were able to extract and quantify 323 proteins (>75% of samples) from single 2 cm-long HS segments. Compared to dark HS collected at the same time from the same individuals, white hairs contained several differentially enriched (upregulated) or depleted (downregulated) proteins (**Figure 1J-K** see **Supplemental Table S1** for complete list) on which we performed GO (Gene Ontology) and KEGG (Kyoto Encyclopedia of Genes and Genomes) enrichment analysis and explored their protein-protein interaction networks (**Supplemental Figure S2**). The protein networks for both downregulated (<0.8-fold, n=23) and upregulated (>1.5-fold, n=67) proteins contain significantly more interactions than expected by chance (P<0.00001, observed vs expected protein-protein interactions). Thus, coherent groups of functionally related proteins are differentially expressed in white hairs, from which two main patterns emerged.

The first main pattern relates to protein biosynthesis and energy metabolism. A large fraction (34.3%) of upregulated proteins in white hairs was related to ribosome function, protein processing, and associated cytoskeletal proteins. Upregulation of the machinery responsible for protein synthesis and amino acid metabolism included the ribosomal protein RPS15A, which is known to localize to mitochondria. Of all upregulated proteins in white hairs, 26.8% were known mitochondrial proteins (MitoCarta2.0 and others)(*43*). These proteins are involved in various aspects of energy metabolism, including substrate transport (carnitine palmitoyltransferase 1A, CPT1A; malonate dehydrogenase 1, MDH1), respiratory chain function (Complex III subunit 1, UQCRC1), and catecholamine homeostasis (Catechol-O-Methyltransferase, COMT). White hairs also contained more proteins involved in glucose (glucose 6-phosphate dehydrogenase, G6PD; phosphoglycerate kinase 1, PGK1) and lipid metabolism located in either the mitochondria or cytoplasm (fatty acid synthase, FASN; acyl-CoA thioesterase 7, ACOT7; mitochondrial trifunctional enzyme subunit beta, HADHB) or in peroxisomes (acyl-CoA acyltransferase 1, ACAA1). The metabolic remodeling in white hairs is consistent with the established role of mitochondria and metabolic regulation of hair growth and maintenance in animal models (*12, 44–46*), and possibly consistent with hair anomalies reported in human patients with mitochondrial disease (*47*). The upregulation of energy metabolism may subserve the likely increased energy demands in depigmented hairs. However, our data and those of others (*40*) implicate the upregulation of specific mitochondrial proteins involved, not necessarily in global energy metabolism, but in specific metabolic activities such as amino acid and lipid biosynthesis.

A second less robust pattern relates more directly to melanosome biology. In line with the lysosomal origin of melanosomes that are largely absent in depigmented HS (*37*), several lysosomal proteins (PLD3, CTSD, HEXB, and LAMP1) were downregulated in white hairs, consistent with previous literature (*40*). White hair shafts also showed a depletion of six main keratins (see **Figure S2**), likely because graying can affect the nature of keratinocytes proliferation (*11*), of proteins associated with exocytosis, such as ITIH4 and APOH (potentially involved in the secretion of melanosomes from melanocytes to keratinocytes), as well as proteins associated with mitochondrial calcium transmembrane transport. Interestingly, calcium exchange between mitochondria and the melanosome is required for melanin pigment production in melanocytes (*14*), and calcium signaling promotes melanin transfer between melanosomes and keratinocytes (*48*).

Finally, canities-affected white HFs also showed upregulation of antioxidant proteins, specifically those localized to mitochondria and cytoplasm (superoxide dismutase 1, SOD1; peroxiredoxin 2, PRDX2), in line with the role of oxidative stress in HS depigmentation (*7, 49*). Alterations among these individual metabolic and mitochondrial enzymes were further reflected in KEGG pathways functional enrichment analyses indicating a significant enrichment of metabolic pathways including carbon metabolism and fatty acid synthesis, amino acids, and oxidative phosphorylation (see below).

### Validation of graying-associated proteomic signatures

To independently validate these results, we extended this analysis to white and dark HS from 6 individuals (3 males, 3 females, range: 24-39 years) analyzed on a separate proteomic platform and in a different laboratory. In this experiment, a total of 192 proteins (≥3 samples) were quantified from 1cm-long HS segments. This dataset showed a similar trend as the first analysis towards a preferential overexpression of proteins with graying (55% upregulated vs 29% downregulated in white HS) (**Figure 1L-M,** see **Supplemental Table S2** for a complete list). The most highly upregulated proteins included mitochondrial components such as the voltage-dependent anion channel 1 (VDAC1), a subunit of ATP synthase (ATP5A1), and a regulator of mitochondrial respiratory chain assembly (Prohibitin-2, PHB2). Again, the antioxidant enzyme SOD1 was enriched in white relative to dark HSs.

To examine the possibility that these relative upregulations are driven by a global downregulation of highly abundant housekeeping proteins, we analyzed the intensity-based absolute quantification (iBAQ) data for each sample. This confirmed that the housekeeping proteins, including keratins and keratin-associated proteins, were not downregulated in white hairs, but generally unchanged or slightly upregulated. Moreover, as a whole, upregulated proteins formed a coherent protein-protein interactions cluster (p<0.00001) and pathway analysis showed a significant enrichment of carbon metabolism, glycolysis/glucogenesis, pyruvate metabolism, and amino acid synthesis pathways in white relative to dark HS (**Supplemental Figure S3, Figure 4E**). On the other hand, proteins downregulated in white HSs were related to cholesterol metabolism, complement-coagulation cascades, and secretory processes shared with immune/inflammatory pathways (**Figure 4E**). The downregulation of secretory pathways is again consistent with reduced transfer of pigmented melanosomes from the melanocytes to the keratinocytes.

To verify the robustness of these findings using an alternative analytical approach, we built a simple partial least square discriminant analysis (PLS-DA) multivariate model, which provided adequate separation of white vs dark HS (**Supplemental Figure S4**). Simple interrogation of this model to extract the features (i.e., proteins) that maximally contribute to group separation yielded a set of proteins enriched for estrogen signaling pathways, complement and coagulation cascades, as well as metabolic pathways including NAD^+^/NADH, cholesterol, pyruvate, and carbon metabolism, similar to results above. Interestingly, we also identified 13 proteins that were undetectable in any of the dark HS (either not expressed, or below the detection limit), but consistently present in white HS (**Supplemental Table S3**). These proteins are either newly induced or experience a substantial upregulation with graying (fold change tending towards infinity). A separate functional enrichment analysis for these induced proteins also showed significant enrichment for the same aging-related metabolic pathways as for the upregulated protein list: glycolysis/glucogenesis, carbon, pyruvate, and cysteine and methionine metabolism (all P<0.001).

These converging proteomics data, which are consistent with previous findings (*40*), support a multifactorial process directly implicating metabolic changes in human hair graying (*6*). Moreover, given that metabolic pathways are rapidly and extensively remodeled by environmental and neuroendocrine factors – i.e., they naturally exhibit plasticity – these data build upon previous proteomic evidence to show that human hair graying could be, at least temporarily, reversible

### Human hair graying is, at least temporarily, reversible

Our analysis of HPPs in healthy unmedicated individuals revealed several occasions whereby white hairs naturally revert to their former dark pigmented state. This phenomenon was previously reported only in a handful of case reports, with only a single two-colored HS in each case(*30*). Here we document the reversal of graying along the same HS in both female and male individuals, ranging from a prepubescent child to adults (age range 9 to 39 years), and across individuals of different ethnic backgrounds (1 Hispanic, 12 Caucasian, 1 Asian). This phenomenon was observed across frontal, temporal, and parietal regions of the scalp (**Figure 2A**), as well as across other corporeal regions, including pubic (**Figure 2B**) and beard hairs (**Figure 2C**). The existence of white HS undergoing repigmentation across ages, sexes, ethnicity, and corporeal regions documents the reversibility of hair graying as a general phenomenon not limited to scalp hairs. Nevertheless, we note that this phenomenon is limited to rare, isolated hair follicles. As their occurrence will probably go unnoticed in most cases, it is difficult to assess the true incidence of these events. Still, only a limited number of case studies reporting natural reversibility of graying appear in the literature, and we were only able to identify 14 participants over an active recruitment period of 2.5 years, indicative of the rarity of this phenomenon.

**Fig. 2.**
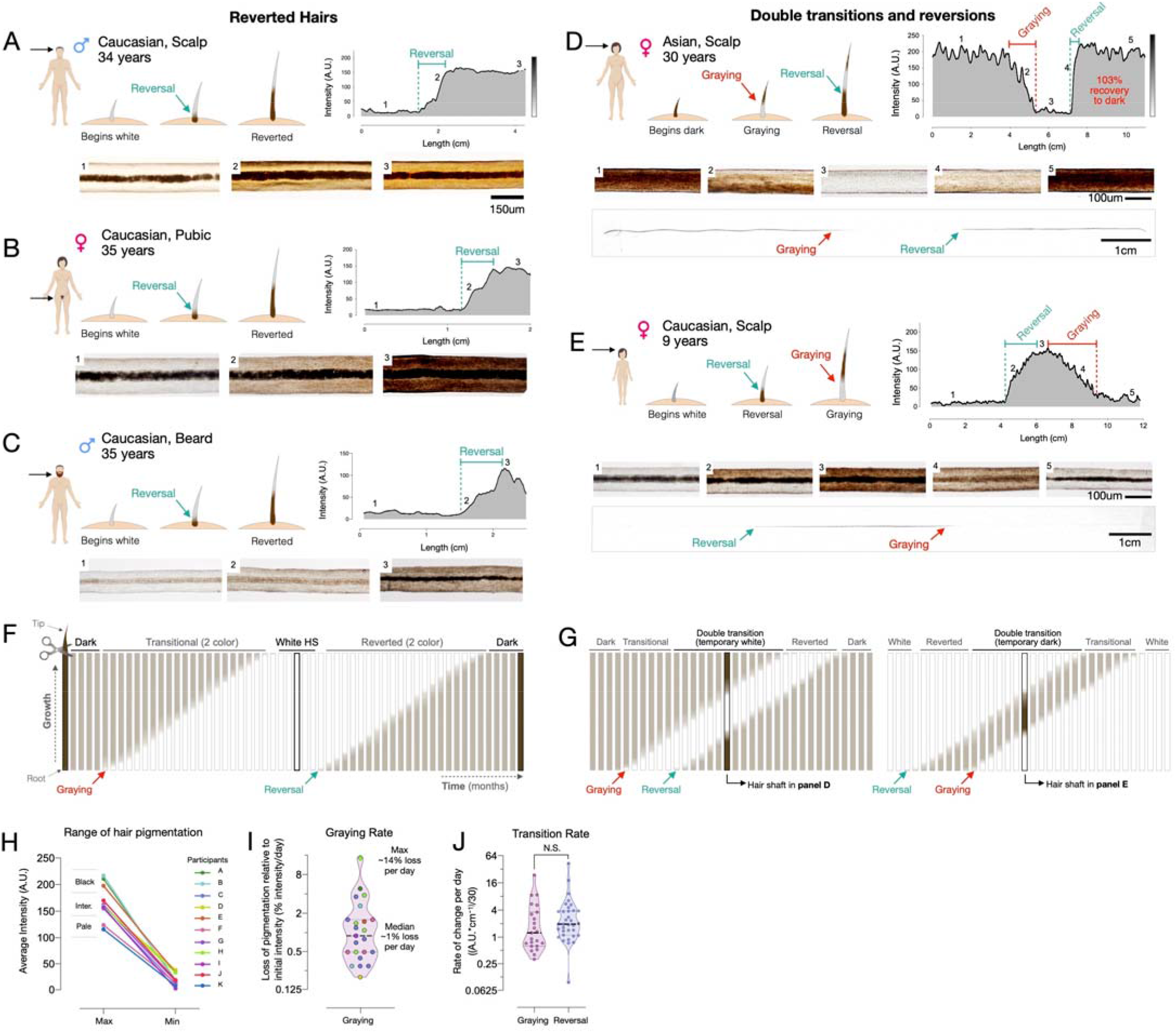
Reversal of hair graying across ages and body regions. (**A-G**) Examples of HS graying and reversal including schematic of hair growth (*top left*), digitized HPP (*top right*), and light microscopy images (*bottom*) corresponding to numbered HS segments on the HPP plot. (**A**) Examples illustrating the reversal of graying along the length of scalp, (**B**) pubic, (**C**) and beard human HSs. (**D**) Example of segmental HS with double transitions, including temporary graying and (**E**) temporary reversal from an adult and a child, respectively. See Supplemental Figure S5 for additional examples and Movie S1 for animation. (**F**) Time course diagram illustrating the progression of a single dark HS undergoing graying followed by reversal back to its original color, and (**G**) closely occurring events of graying and reversal occurring, producing HS with double transitions. (**H**) Average maximum and minimum pigmentation intensity among transitioning hairs from participants with two-colored hairs (n=11). Hairs with an average maximum intensity >180 A.U. are categorized as high intensity (black), 140-180 A.U. as intermediate intensity, and 100-140 A.U. as low intensity (pale color), indicating that these findings generalize across a range of pigmentation densities. (**I**) Rate of depigmentation per day in graying HS (n=23), measured from the slope on HPP graphs expressed as % of starting intensity loss per day (assuming growth rate of 1cm/month). (**J**) Comparison of the absolute rate of pigmentation change per day in graying (n=23) and reverted (n=34) HS. I and J are reported on a log_2_ scale to facilitate visualization.

Moreover, more complex HPPs with double transitions and reversions in the same HS were observed in both directions: HS undergoing graying followed by rapid reversal (**Figure 2D**), and repigmentation rapidly followed by graying (**Figure 2E**). Importantly, both patterns must have occurred over the course of a single anagen (growth) phase in the hair growth cycle, implicating cellular mechanisms within the HFPU. Greatly extending previous case studies of these rare hair repigmentation events, the current study provides the first quantitative account of the natural and transitory reversibility of hair graying in humans.

We understand the emergence of a reverted HS – that is, a HS with a white distal segment but with a dark proximal end – as necessarily having undergone repigmentation to its original pigmented state following a period of time as a depigmented “old” white hair (**Figure 2F**). In double transition HS with three segments, repigmentation must take place within weeks to months after graying has occurred, producing three distinct segments present on the same hair strand (**Figure 2G**). Microscopic imaging along the length of a single HS undergoing a double transition (graying followed by rapid reversal) can be visualized in **Movie S1**, illustrating the dynamic loss and return of pigmented melanosomes within the same HS.

Our hair digitization approach also provides the first estimates of the rates of change in pigmentation for HS covering a broad range of initial colors and darkness (**Figure 2H**). Across individuals, assuming a scalp hair growth rate of 1 cm/month (*34*), the rates of depigmentation in graying hairs ranged widely from 0.3 to 23.5 units of hair optical density (scale of 0-255 units) per day, corresponding to between 0.2% and 14.4% loss of hair color per day (**Figure 2I**). The rate at which HS regain pigmentation during reversal was 0.1 to 42.5 units per day, which is similar (~30% faster on average) to the rate of graying (Cohen’s d = 0.15, p = 0.59) (**Figure 2J**). Given these rates, the fastest measured transitioning hairs gray and undergo full reversal in ~3-7 days (median: ~3 months). Thus, rather than drifting back towards the original color, repigmentation of white human HS occurs within the same time frame and at least as rapidly as the process of graying itself.

The spectrum of graying transitions and reversals patterns observed in our cohort, including measured rates of repigmentation along individual hairs, is shown in **Supplemental Figure S5**. These results establish the wide range of naturally-occurring rates of pigmentary changes in single hairs, which vary by up to an order of magnitude from hair to hair. These data also suggest that reversal/repigmentation may reflect the action of as yet unknown local or systemic factors acting on the HFPU within a time frame of days to weeks.

### Correlated behavior of multiple scalp hairs

We then asked whether the reversal of graying is governed by a process that is unique to each human scalp HF or if it is likely coordinated by systemic factors that simultaneously affect multiple HFs. Participants’ scalps were visually inspected to identify two-colored hairs, including both graying transitions and reversal. In our combined cohort, three individuals had multiple two-colored hairs collected at either one or two collection times within a one-month interval. In each case, the multiple two-colored hairs originated from independent HFs separated by at least several centimeters (e.g., left vs right temporal, or frontal vs temporal). If the hairs are independent from each other, the null hypothesis is that different HSs will exhibit either graying or reversal changes and will have independent HPPs. If multiple HSs were coordinated by some systemic factor, then we expect HPPs to exhibit similarities.

In a first 35-year-old female participant with dark brown hair, 3 two-colored hairs were identified at a single instance of collection. Notably, all three hairs exhibited dark-to-white graying. Moreover, when the HPPs of the 3 hairs were quantified and plotted, the HPPs followed strikingly similar graying trajectories (r=0.76-0.82) marked by a similar onset of graying, similar HPP intermittent fluctuations (note the rise ~10 cm), and a similar time point where all hairs become fully depigmented (~15 cm) (**Figure 3A**). A permutation test on the similarity of the color trajectories yielded p=0.032, suggesting possible synchrony between different HSs. If our simulation considers only hairs that transition in one direction (from dark to white) this gives p=0.086 (see *Methods* for details).

**Figure 3.**
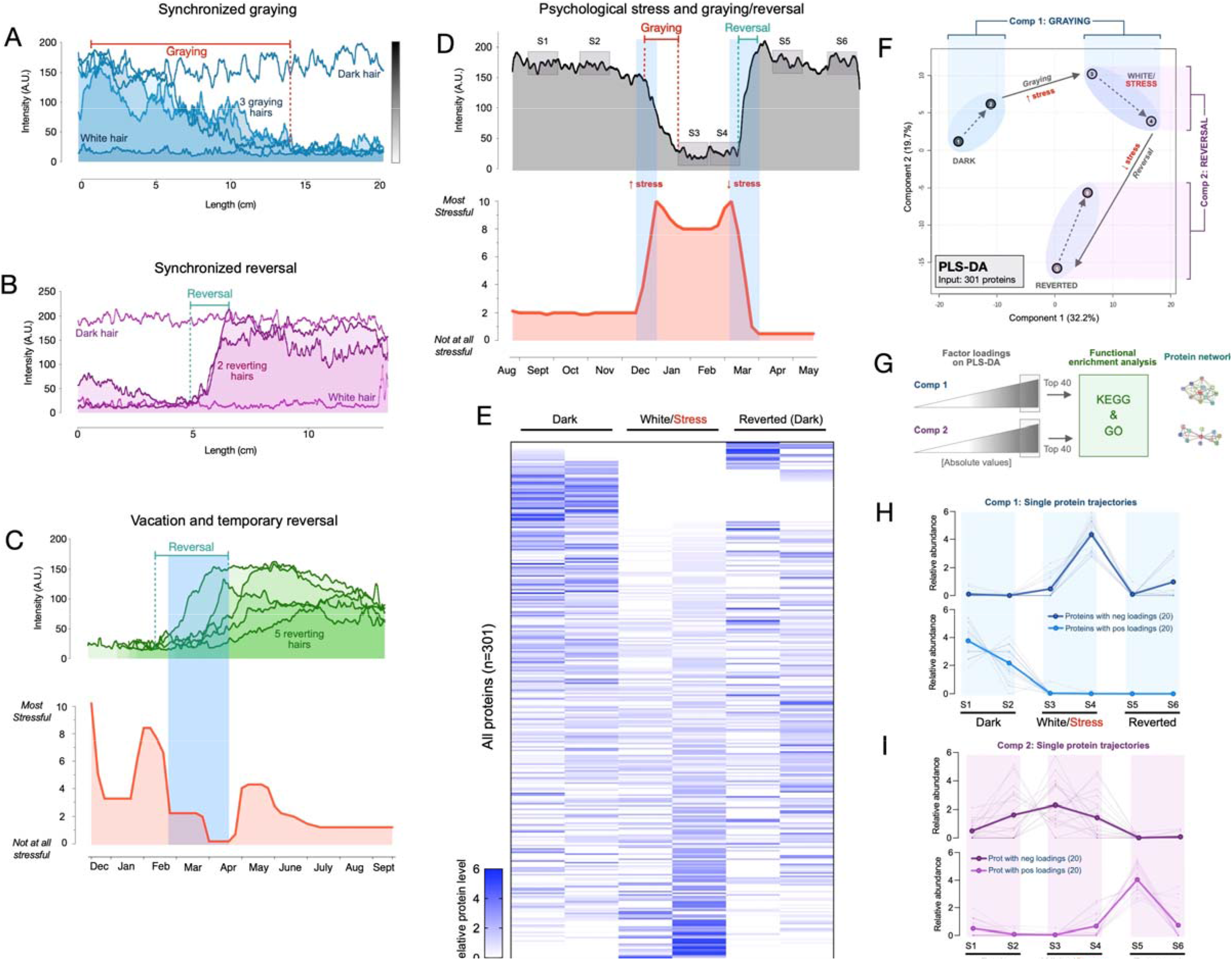
Synchronous graying and reversal behavior across multiple hairs and associations with psychosocial stress. (**A**) In a 35-year-old Caucasian female, multiple HS (n=3) undergoing graying simultaneously. (**B**) In a 37-year-old Caucasian female, two bi-color HS collected two months apart aligned based on a growth rate of 1 cm/month undergoing reversal nearly simultaneously. In A and B, simultaneously plucked dark and white hairs are plotted for reference. (**C**) In a 35-year old Caucasian male, multiple bi-color HS (n=5) undergoing reversal (top-panel) plotted against time-matched self-reported psychosocial stress levels (bottom-panel). (**D**) HS from a 30-year old Asian female with two months of self-reported profound perceived stress associated with temporary hair graying and reversal. Note the synchronicity between the increase in stress and rapid depigmentation (i.e., graying), followed by complete (103%) recovery of HS pigmentation (i.e., reversal of graying) upon alleviation of life stress. (**E**) Heatmap of protein abundance (n=301) across 6 segments: 2 dark prior to stress/graying, 2 white following graying, 2 dark segments after reversal. (**F**) Multivariate PLS-DA model of the 6 segments from the HS in E, highlighting the model’s first and second principal components related to graying and reversal, respectively. Numbers 1 to 6 correspond to HS segments on D. (**G**) Factor loadings for Components 1 and 2 were used to extract the most significant proteins, subsequently analyzed for functional enrichment categories in KEGG and GO databases, and STRING protein networks. (**H**) Trajectories of protein abundance from the top Component 1 and (**I**) Component 2 features across the 6 segments; proteins with positive (*top*) and negative loadings (*bottom*) are shown separately.

**Figure 4.**
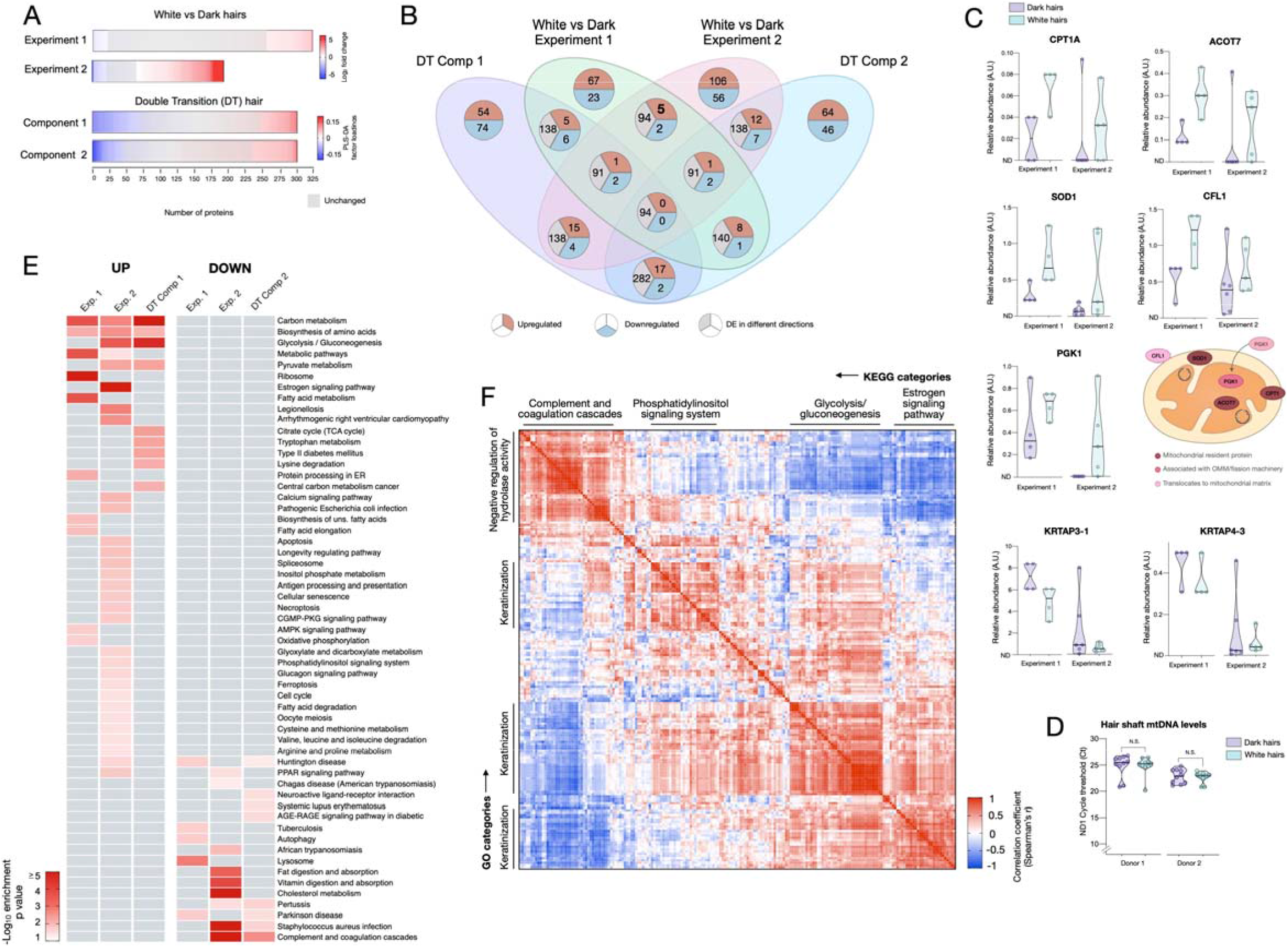
Meta-analysis of human hair proteomic findings comparing dark and white hairs. (**A**) Number of downregulated (<0.8-fold, blue) and upregulated (<1.5-fold) proteins across datasets, and unchanged proteins shown in grey. (**B**) Venn diagram illustrating the intersection of datasets. The number of overlapping proteins across datasets that are either consistently down- or upregulated, or proteins not regulated in the same direction, are shown for each area of overlap. (**C**) Individual protein abundance for consistently upregulated (n=5) and downregulated proteins (n=2) across experiments 1 and 2 shown are shown as box and whiskers plots, with bars extending from the 25^th^-75^th^ percentiles, and whiskers from min to max values. Lines indicate the median and “+” signs indicate the mean. Fold difference values are the mean fold differences relative to dark hairs. (**D**) Mitochondrial DNA abundance in human HS of the same two donors as in Figures 1F-I (AA male, Cau male). (**E**) Summary of significantly enriched KEGG categories across datasets, for upregulated (*left*) and downregulated (*right*) proteins. (**F**) Correlation matrix (Spearman’s r) of all detected proteins (n=192) in experiment 2, illustrating general human hair protein co-expression across dark and white pigmented states (dark, white). Four main clusters are highlighted and labeled by their top KEGG category. N.S. from Mann Whitney Test.

In a 37-year-old female participant with brown hair, two transition hairs were identified. The HPPs for both hairs revealed strikingly similar trajectories (r=0.80), in this case undergoing spontaneous reversal in a near-synchronous manner upon alignment (p<0.001 when considering hairs transitioning in either direction, and similarly, p<0.001 considering only hairs transitioning from white to dark, **Figure 3B**). Thus, these findings extend previous reports in single isolated hairs by providing quantitative accounts of coordinated HS (re)pigmentation across multiple hairs.

Candidate humoral hair pigmentation modulators that could create synchrony in graying or repigmenting hairs include neuropeptides, redox balance, and steroid or catecholamine hormones (*25, 33, 50, 51*) that can directly regulate the human HFPU (*6*), impact intrafollicular clock activity (*50*), or regulate the expression of other melanotropic neurohormones in the human HFPU such as α-MSH, ß-endorphin, and TRH (*52*). These factors must act in parallel with genetic factors that influence inter-individual differences in aging trajectories.

### Hair graying and reversal are linked to psychosocial stress levels

In light of these results, we next applied our HPP method to examine the possibility that psychological stress is associated with hair graying/reversal in humans. Anecdotal case reports suggest that psychological stress and other behavioral factors accelerate the hair graying process (*53*), a notion recently supported by studies in mice demonstrating that adrenergic stimulation by norepinephrine signaling leads to melanocyte stem cell depletion in mice (*33*). However, contrary to mice where this process appears to be irreversible at the single hair follicle level, our data demonstrates that human hair graying is, at least under some circumstances, reversible. This dichotomy highlights a fundamental difference between rodent and human HF biology, calling for a quantitative examination of this process in humans.

As evidence that environmental or behavioral factors influence human hair graying, epidemiological data suggests that smoking and greater perceived life stress, among other factors, are associated with premature graying (*2*). Chronic psychosocial stress also precipitates telomere shortening, DNA methylation-based epigenetic age, as well as other biological age indicators in humans (*32, 54*), demonstrating that psychological factors can measurably influence human aging biology. In relation to mitochondrial recalibrations, psychosocial factors and induced stress can also influence mitochondrial energetics within days in humans (*55*) and animals (*56*). To generate proof-of-concept evidence and test the hypothesis that psychosocial or behavioral factors may influence graying at the single-HF level, we leveraged the fact that HPPs reflect longitudinal records of growth over time – similar to tree rings – which can be aligned with assessments of life stress exposures over the past year. By converting units of hair length into time, perceived stress levels can be quantitatively mapped onto HPPs in both graying and transitional hairs.

A systematic survey of two-colored hairs on the scalp of a 35-year-old Caucasian male with auburn hair color over a two-day period yielded five two-colored HS from the frontal and temporal scalp regions. Again, two-colored hairs could either exhibit depigmentation or reversal. Unexpectedly, all HS exhibited reversal. HPP analysis further showed that all HS underwent reversal of graying around the same time period. We therefore hypothesized that the onset of the reversal would coincide with a decrease in perceived life stress. A retrospective assessment of psychosocial stress levels using a time-anchored visual analog scale (participants rate and link specific life events with start and end dates, see *Methods* and **Supplemental Figure S6** for details) was then compared to the HPPs. The reversal of graying for all hairs coincided closely with the decline in stress and a 1-month period of lowest stress over the past year (0 on a scale of 0-10) following a two-week vacation (**Figure 3C**).

We were also able to examine a two-colored hair characterized by an unusual pattern of complete HS graying followed by rapid and complete reversal (same as in Figure 2B) plucked from the scalp of a 30-year-old Asian female participant with black hair. HPP analysis of this HS showed a white segment representing approximately 2 cm. We therefore hypothesized that this reversible graying event would coincide with a temporary increase in life stress over the corresponding period. Strikingly, the quantitative life stress assessment over the last year revealed a specific 2-month period associated with an objective life stressor (marital conflict and separation, concluded with relocation) where the participant rated her perceived stress as the highest (9-10 out of 10) over the past year. The increase in stress corresponded in time with the complete but reversible hair graying (**Figure 3D**). This association was highly significant (p=0.007) based on our permutation test. Given the low statistical probability that these events are related by chance, life stress is the likely preceding cause of these HS graying and reversal dynamics. These data demonstrate how the HPP-stress mapping approach makes it possible to examine the coordinated behavior of graying and reversal dynamics with psychosocial factors, raising the possibility that systemic biobehavioral factors may influence multiple HFs simultaneously and regulate HPPs among sensitive hairs.

### Single-hair longitudinal proteomic signature

To assess whether rapid graying and reversal events among a single hair are molecularly similar or distinct to those described in the two proteomics experiments above, we dissected 6 segments (2 dark, 2 white, 2 reverted) of the single HS in Figure 3D and quantified their proteomes as part of Experiment 2. This produced a longitudinal, single-hair, proteomic signature (**Figure 3E**) containing 301 proteins quantified in ≥2 of the 6 segments. To examine how the proteome as a whole is altered through the graying and reversal transitions associated with psychosocial stress levels, we generated a PLS-DA model with all 6 segments. Both dark segments clustered together, with similar values on both first and second principal components. The white and reverted segments clustered in separate topological spaces (**Figure 3F**). Graying was associated with a positive shift largely along the first component (Component 1), whereas reversal was associated with a negative shift on the second component (Component 2) and a more modest negative shift in Component 1. We therefore extracted loading weights of each protein on Components 1 and 2 (reflecting each protein’s contribution to group separation) and used the top proteins (n=20 highest negative and 20 most positive loadings, total n=40 per component) to interrogate KEGG and GO databases.

The model’s Component 1 (associated with graying) contained proteins that were either i) not expressed in the dark HS but induced selectively in the white HS segment, or ii) highly abundant in dark segments but strongly downregulated in white and reverted segments (**Figure 3H**, *top* and *bottom*, respectively). In gene set enrichment analysis of Component 1 (graying), the top three functional categories were carbon metabolism, glycolysis/gluconeogenesis, and Kreb’s cycle (**Figure 4E**). Component 2 (reversal)-associated proteins exhibited distinct trajectories either peaking in the first white segment or upon reversal (**Figure 3I**) and mapped to pathways related to the complement activation cascade, infectious processes, and Parkinson’s and Huntington’s disease (**Figure 4E**). In contrast, a null set of hair proteins not contributing to either components exhibited enrichment for extracellular exosomes and cell-cell adhesion that reflect hair shaft biology (**Supplemental Figure S7**), therefore illustrating the specificity of our findings related to graying and reversal. These data indicate that the reversal of graying at the single-hair level is not associated with a complete reversal in the molecular composition of the HS. Rather, some of the proteomic changes in hair graying are enduring despite successful repigmentation.

### Conserved proteomic signatures of hair graying

To systematically examine the overlap among the different proteomic datasets and to derive functional insight into the hair graying process in humans, we then integrated results from the three datasets described above. White HS show consistently more upregulated than downregulated proteins across datasets (2.91-fold in Experiment 1, 1.89-fold in Experiment 2) (**Figure 4A**). This preferential upregulation suggests that the depigmentation process likely involves active metabolic remodeling rather than a passive loss of some pigmentation-related factor. The overlap in the specific proteins identified across dark-white comparisons and among the 6-segments hair is illustrated in **Figure 4B**.

Five proteins were consistently upregulated between experiments 1 and 2. These include three well-defined resident mitochondrial proteins involved in lipid metabolism: CPT1A, which imports fatty acids into mitochondria (*57*); ACOT7, which hydrolyzes long-chain fatty acyl-CoA esters in the mitochondrial matrix and cytoplasm (*58*); and SOD1, which dismutates superoxide anion into hydrogen peroxide (H_2_O_2_) in the mitochondrial intermembrane space (*59*). The other two proteins include the actin-depolymerizing protein cofilin-1 (CFL1) and the core glycolysis enzyme phosphoglycerate kinase 1 (PGK1) (**Figure 4C**). Interestingly, CFL1 promotes mitochondrial apoptotic signaling via cytochrome c release (*60*) and regulates mitochondrial morphology via its effect on actin polymerization-dependent mitochondrial fission (*61*). And although PGK1 is a cytoplasmic kinase, it was recently demonstrated to translocate inside mitochondria where it phosphorylates and inhibits pyruvate dehydrogenase and Krebs cycle activity (*62*). Thus, all five proteins validated across both experiments are linked to mitochondrial energy metabolism, implicating mitochondrial remodeling as a feature of hair graying. Interestingly, all five proteins have also been linked to the biology of melanocytes (*63–67*), the source of pigment in the HFPU. The downregulated proteins were keratins, with small effect sizes, and not particularly robust. Analysis of the intensity based on absolute quantification (iBAQ) data confirmed the upregulation of these five mitochondrial proteins, and the absence of substantial changes in the keratins. Together, these data suggest that HS proteome profiling may provide a retrospective access to some aspect of melanocyte metabolism, which opens new possibilities to study HF aging biology.

Since the observed proteomic signatures are related to specific metabolic pathways rather than the typical high-abundance mitochondrial housekeeping proteins, we reasoned that the upregulation of these mitochondrial components unlikely reflects a bulk increase in total mitochondrial content. To investigate this point using an independent method, we quantified mitochondrial DNA (mtDNA) abundance in human HS by real-time qPCR. Both white and dark HSs contain similarly high levels of mtDNA (**Figure 4D**). The same was true in the follicles of the same hairs (**Supplemental Figure S8**). The similar mtDNA levels between dark and white hairs increases the likelihood that the reported proteomic changes reflect the induction of specific metabolic pathways associated with hair graying rather than bulk increase in mitochondrial mass.

Finally, to identify a general proteomic signature of graying hair, we compiled the enrichment scores for KEGG pathways across all datasets (**Figure 4E**). Consistent with the function of the individual proteins identified in both group comparisons (Experiments 1 and 2) and the multi-segment double-transition hair, white HS showed consistent upregulation of carbon metabolism and amino acid biosynthesis, glycolysis/gluconeogenesis, and general metabolic pathways relevant to aging biology (*68*). Comparatively fewer pathways were consistently represented for downregulated proteins across independent experiments. In relation to hair biology, our data adds to previous efforts (*40*) and provides a quantitative map of the co-expression among keratin and non-keratin HS proteins across dark and white hairs (**Figure 4F**). Computing the cross-correlations for each protein pair revealed four main clusters among the HS proteome. As expected for hair, keratins were well-represented and constituted the main GO biological processes category for 3 of the 4 clusters. The top KEGG categories included glycolysis and estrogen signaling pathways, which also showed strong co-expression with each other, highlighting potential interaction among endocrino-metabolic processes in relation to human hair pigmentation. In general, the identification of several non-keratin metabolism-related proteins in the HS opens new opportunities to investigate graying pathobiology and to non-invasively access past molecular and metabolic changes that have occurred in the aging HFPU of the dynamically growing hair.

### In silico modeling of hair graying and its temporary reversal

Finally, to narrow the range of plausible mechanisms for the observed age-related graying and reversal events, we developed a simulation model of HPPs. Graying dynamics of an individual’s hair population (~100,000 hairs) across the average 80-year lifespan cannot practically be measured. In the absence of such data, we propose here a mathematical model to simulate hair graying trajectories across the human lifespan (**Figure 5A**, available online, see *Methods* for details) as has been attempted previously for hair growth cycles (*69*). As basic tenets for this model, it is established that *i)* the onset of human hair graying is not yet underway and rarely begins in childhood, *ii)* graying routinely starts between 20-50 years of age, *iii)* graying is progressive in nature (the total number and percentage of gray hairs increases over time), and *iv)* the proportion of white hairs reaches high levels in old age, although some hairs can retain pigmentation until death, particularly among certain body regions (*8*). Additionally, our findings demonstrate that *v)* age-related graying is naturally reversible in isolated hair follicles, at least temporarily and in individual HS, and may be acutely triggered by stressful life experiences, the removal of which can trigger reversal.

**Figure 5.**
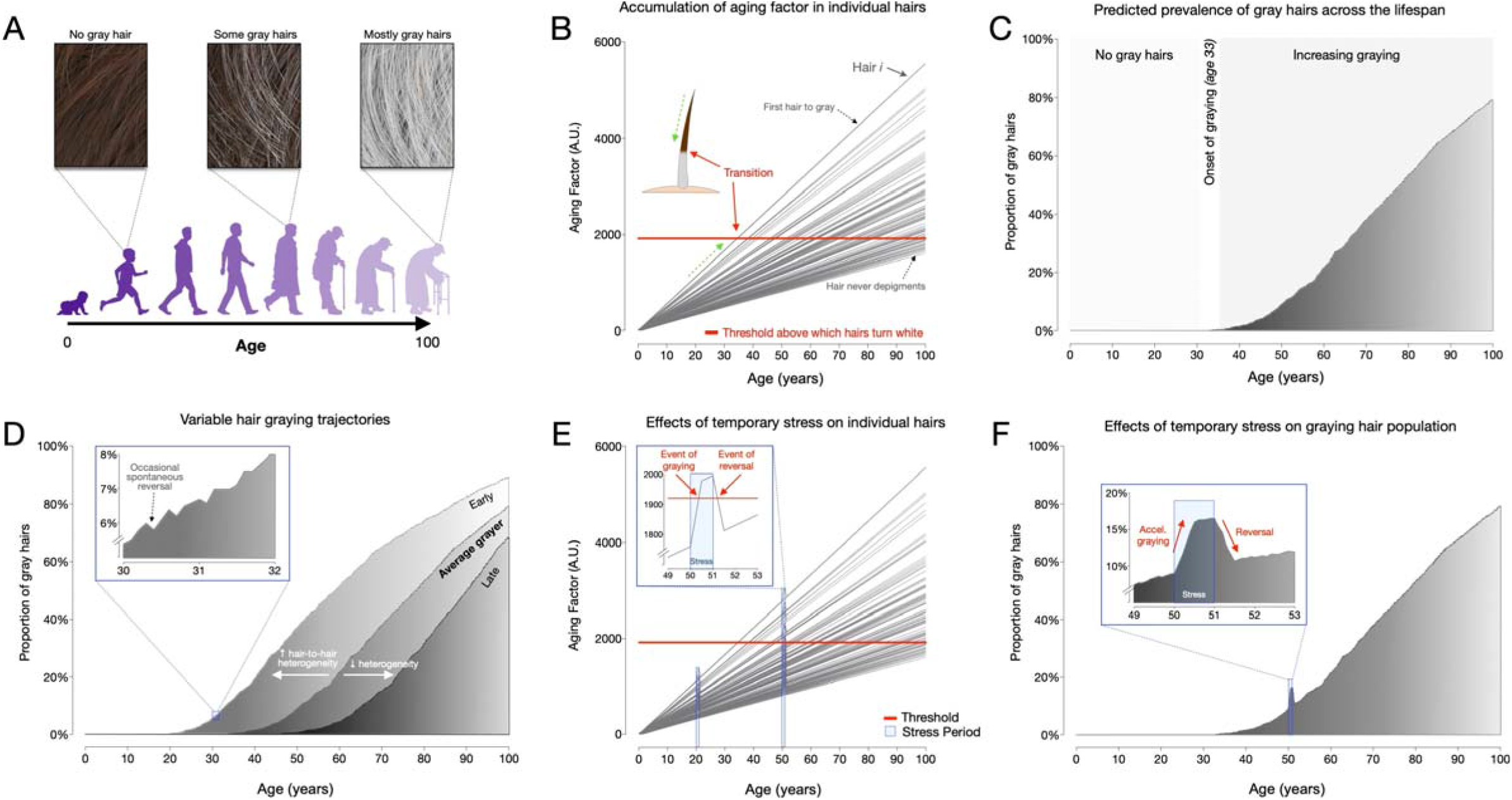
Modeling of hair graying and reversal across the human lifespan and in response to temporary stress. (**A**) Schematic overview of the average graying process across the lifespan involving the gradual loss of pigmentation, or accumulation of white hairs, mostly in the second two-thirds of life. (**B**) Depiction of individual hairs (each line is a hair, i) from a linear mixed effects model with random intercept and slopes predicting hair graying. The model assumes i) a constant increase in a putative aging factor and ii) a constant threshold above which hairs undergo depigmentation. All model parameters are listed in Supplemental Table S4. (**C**) Predicted hair population behavior (n=1,000 hairs) shown as a cumulative frequency distribution of white hairs from the individual trajectories in panel (B). (**D**) Frequency distributions of gray hairs for individuals with early (*left)*, average (*middle*), or late (*right*) hair graying phenotypes. The inset highlights a 2-year period during mid-life indicating gradual accumulation of white hairs, as well as the spontaneous repigmentation of a small fraction of white hairs (decrease in % white hairs), consistent with real-world data. **(E**) Single hair-level and (**F**) hair population-level results from the addition of two acute stress periods (each one year in duration, occurring at ages 20 and 50). The optimized model accounts for stress-induced graying in hairs whose aging factor is close to the depigmentation threshold, but not for young hairs or those far away from the threshold. Similarly, the removal of the stressor causes repigmentation of hairs when the aging factor returns below the threshold.

Aiming for the simplest model that accounts for these known features of hair graying dynamics, we found a satisfactory model with three components (**Figure 5B**): 1) an “*aging factor*” that progressively accumulates within each hair follicle, based on the fact that biological aging is more accurately modeled with the accumulation of damage, rather than a decline in stem cells or other reserves (*70*); 2) a biological *threshold*, beyond which hairs undergo depigmentation (i.e., graying), characterizing the transition between the dark and white states in the same HS; and 3) a “*stress factor*” that acutely but reversibly increases the aging factor during a stressful event. For modeling purposes, the accumulation of the aging factor is equivalent to the inverse of the decrease in a youth factor (e.g., loss of telomere length with age). Based on the mosaic nature of scalp HFs and our data indicating that not all hairs are in perfect synchrony, the proposed model for an entire population of hairs must also allow a variety of aging rates, as well as differential sensitivity to stress among individual hairs.

We find that the model’s predicted hair population behavior (% of white HSs on a person’s head over time) across the lifespan is consistent with expected normal human hair graying dynamics (**Figure 5C**). White hairs are largely absent until the onset of graying past 20 years of age then accumulate before finally reaching a plateau around 70-90% of white hairs, near 100 years. Thus, this model recapitulates the expected between-hair heterogeneity of graying within an individual, producing the common admixture of white and pigmented hairs or “salt & pepper” phenotype in middle-age. However, some individuals also develop hairs with intermediate pigmentation states (i.e., silver/steel color), which our model does not reproduce. This represents a limitation to be addressed in future research.

We note that there are natural inter-individual differences in the rate of graying: some individuals begin graying early (onset in early 20’s); some begin late (onset in 50’s). A higher rate of accumulation of the aging factor (higher slope for each hair) or a lower threshold naturally accounts for earlier onset of graying. In addition, our model reveals that within a person, greater hair-to-hair heterogeneity in the rate of aging between HFs, modeled as the standard deviation of slope across hairs, also influences the onset of graying. Greater heterogeneity between HFs allows for earlier onset of graying, whereas decreasing hair-to-hair variation (i.e., lower heterogeneity) is associated with a “youthful” later onset of graying (**Figure 5D**). Interestingly, this unpredicted result aligns with the notion that increased cell-to-cell heterogeneity is a conserved feature of aging (*71–73*) and that biological heterogeneity can predict all-cause mortality in humans (*74*).

### Modeling stressors produce hair graying and temporary reversal

Using parameter values that yield the average onset and rate of graying, we then simulated the influence of acute psychosocial stressors, either early in life before the onset of graying, or later once gray HSs have begun to accumulate. Similar to our data, the model also predicts transitory, or temporary reversible events of graying (see **Figure 3D**). Transitory graying events do not affect all hairs, only those that are close to the threshold at the time of stress exposure undergo graying. Hairs whose cumulative aging factors are substantially lower than threshold do not show stress-induced graying (a 5-year-old is unlikely to get gray hairs from stress, but a 30-year-old can) (**Figure 5E-F**). Similarly, gray hairs far above threshold are not affected by periods of psychosocial stress. Thus, our model accounts for both the overall hair graying dynamics across the lifespan, and how a stressor (or its removal) may precipitate (or cause reversal of) graying in hairs whose aging factor is close to the graying threshold.

We speculate that this simulation opens an attractive possibility whereby HPP data from individuals could be used in models to formulate predictions about future graying trajectories, and to use HPPs and hair population graying behavior to track the effectiveness of behavioral and/or therapeutic interventions aimed at modifying human aging. Extending our high-resolution quantitative digitization approach to hundreds of randomly sampled dark non-transitioning hairs from different scalp regions in the same individuals, we also show that fully dark (i.e., non-graying) HSs exhibit mostly unique HPPs, but that hairs among the same scalp regions may exhibit more similar HPPs than hairs sampled from different regions (**Figure S9**) (*75*). This may in part be influenced by the migration of stem cells during embryogenesis to different parts of the scalp, or by other unknown factors. This preliminary extension of the HPP methodology provides a foundation for future studies. Moreover, the regional segregation of HPPs may reflect well-recognized regional differences in the rate of HS formation (*76*). Thus, future models may also be able to leverage information contained within HPPs from non-graying hairs and make specific inference from hairs collected across scalp regions. Similar to how decoding temporal patterns of electroencephalography (EEG) provides information about the state of the brain, our data make it imaginable that decoding HPP analysis over time may provide information about the psychobiological state of the individual.

## Discussion

Our approach to quantify HPPs demonstrates rapid graying transitions and their natural transitory reversal within individual human hair follicles at a higher frequency and with different kinetics than had previously been appreciated. The literature generally assumes pigment production in the HFPU to be a continuous process for the entire duration of an anagen cycle, but here we document a complete switch-on/off phenomena during a single anagen cycle. The proteomic features of hair graying directly implicate metabolic pathways that are both reversible in nature and sensitive to stress-related neuroendocrine factors. This result therefore provides a plausible biological basis for the rapid reversibility of graying and its association with psychological factors, and also supports the possibility that this process could be targeted pharmacologically. Melanogenesis is also known to both involve and respond to oxidative stress, a byproduct of mitochondrial metabolic processes (*77*) and driver of senescence (*78*). Moreover, alterations in energy metabolism is a major contributor to other disease-related aging features (*79*), including lifespan regulation (*80, 81*). The upregulation of specific components related to mitochondrial energy metabolism in white hairs suggests that energy metabolism regulates not only hair growth as previously demonstrated (*12, 82, 83*) but also HS graying biology.

Although surprising, the reversal of hair graying is not an isolated case of “rejuvenation” in mammals. *In vivo*, exposing aged mice to young blood in parabiosis experiments (*84, 85*) or diluting age-related factors in old animals (*86*) triggers the reversal of age-related molecular, structural and functional impairments. In human cells, quantitative biological age indicators such as telomere length (*87*) and DNA methylation (*88*) also exhibit temporary reversal in response to exercise and dietary interventions. Moreover, the reversibility of graying in aging human HFs demonstrated by our data is also consistent with the observed reversibility of human skin aging *in vivo* when aged human skin is xenotransplanted onto young nude mice (*89*). Notably this skin “rejuvenation” is associated with a marked increase in the number of melanocytes in human epidermis (*90*), suggesting plasticity of the melanocyte compartment. Therefore, our HPP data and simulation model adds to a growing body of evidence demonstrating that human aging is not a linear, fixed biological process but may, at least in part, be halted or even temporarily reversed. Our method to map the rapid (weeks to months) and natural reversibility of human hair graying may thus provide a powerful model to explore the malleability of human aging biology within time scales substantially smaller than the entire lifespan.

A surprising finding from both proteomics experiments is the bias towards *up*regulation rather than the loss of proteins in depigmented gray HS. As noted above, this may reflect the fact that hair graying is an actively regulated process within the HPFU, and that aging is not marked by a loss, but rather an increase in heterogeneity and biological complexity (*71–73, 91*). Relative to the youthful state, quiescent and senescent cells exhibit upregulation of various secreted factors (*92*), as well as elevated metabolic activities (*93*), rather than global downregulation of cellular activities. Moreover, similar to the macroscopic appearance of hair graying, age-related senescence naturally occurs among cells heterogeneously scattered within tissues (*94*), beginning as early as a few weeks after birth in mice (*95*), and the age-related accumulation of senescence-associated DNA methylation changes among cell populations also exhibit stochasticity (*96*). Our data reveal that the conserved principle of an age-related increase in molecular and cellular heterogeneity is reflected not only at the tissue level (mixture of dark and white hairs) but also in the graying hair proteome. Further work is required to determine if stochastic molecular aging processes, in specific cell types within the HF, account for the macroscopic instability of HFs graying visible on the human scalp.

Finally, in relation to psychobiological processes, the spatio-temporal resolution of the HPP approach provides investigators with an instructive new research tool that allows to link hair graying and reversal events with psychosocial exposures with an unprecedented level of resolution. Here we provided proof-of-concept evidence that biobehavioral factors are linked to human hair graying dynamics. Our optical digitization approach thus extends previous attempts to extract temporal information from human hairs (*97*) and illustrate the utility of HPP profiling as an instructive and sensitive psychobiology research model. Visualizing and retrospectively quantifying the association of life exposures and HPPs may thus contribute to understanding the embedding of stress and other life exposures on human biology.

## Acknowledgements

The authors are grateful to Avsar Rana, David Sulzer, Marko Jovanonic, and to participants who donated hairs and time for this study.

## Author contributions

A.R., S.R., R.K.S., P.P., G.S.: collected data. A.R., S.R., C.L., R.T.O., G.S., M.P.: analyzed data. J.R., R.T.O.: developed the simulation model with S.R., A.R., E.M.: performed imaging. M.P., R.P.: drafted manuscript. A.R. and D.J.T. revised manuscript. M.P., G.S.: designed study. All authors contributed to data interpretation and the final version of the manuscript.

## Data and materials availability

The datasets generated during and/or analyzed during the current study, including electron microscopy analysis (Figures 1, S1), HPPs (Figures 1–3, S5, S9), proteomic data (Figures 1, 3, 4, S2-4, S7, S10), and mtDNA data (Figure 4, S8) are available from the corresponding author upon request. Source code for the hair simulation model is available on the App and on GitHub at https://github.com/junting-ren/hair_simulation. Correspondence and requests for materials should be addressed to Martin Picard

## Supplementary Materials

Materials and Methods Tables S1-S4

Figures S1-S12

References (*98–115*)

Movies S1

## References and Notes

1. D. J. Tobin, The cell biology of human hair follicle pigmentation. Pigment cell & melanoma research 24, 75–88 (2011).

2. A. Akin Belli, F. Etgu, S. Ozbas Gok, B. Kara, G. Dogan, Risk Factors for Premature Hair Graying in Young Turkish Adults. Pediatr Dermatol 33, 438–442 (2016).

3. B. A. Bernard, The human hair follicle, a bistable organ? Experimental Dermatology 21, 401–403 (2012).

4. S. Panhard, I. Lozano, G. Loussouarn, Greying of the human hair: a worldwide survey, revisiting the ‘50’ rule of thumb. Br J Dermatol 167, 865–873 (2012).

5. E. K. Nishimura, S. R. Granter, D. E. Fisher, Mechanisms of hair graying: incomplete melanocyte stem cell maintenance in the niche. Science 307, 720–724 (2005).

6. R. Paus, A neuroendocrinological perspective on human hair follicle pigmentation. Pigment cell & melanoma research 24, 89–106 (2011).

7. P. C. Arck et al., Towards a “free radical theory of graying”: melanocyte apoptosis in the aging human hair follicle is an indicator of oxidative stress induced tissue damage. FASEB J 20, 1567–1569 (2006).

8. R. M. Trueb, D. J. Tobin, Aging Hair. (Springer, Verlag Berlin Heidelberg, 2010).

9. D. J. Tobin, Age-related hair pigment loss. Curr Probl Dermatol 47, 128–138 (2015).

10. W. Nagl, Different growth rates of pigmented and white hair in the beard: differentiation vs. proliferation? Br J Dermatol 132, 94–97 (1995).

11. D. Van Neste, D. J. Tobin, Hair cycle and hair pigmentation: dynamic interactions and changes associated with aging. Micron 35, 193–200 (2004).

12. A. Flores et al., Lactate dehydrogenase activity drives hair follicle stem cell activation. Nat Cell Biol 19, 1017–1026 (2017).

13. R. Williams, M. P. Philpott, T. Kealey, Metabolism of freshly isolated human hair follicles capable of hair elongation: a glutaminolytic, aerobic glycolytic tissue. J Invest Dermatol 100, 834–840 (1993).

14. Z. Zhang, J. Gong, E. V. Sviderskaya, A. Wei, W. Li, Mitochondrial NCKX5 regulates melanosomal biogenesis and pigment production. J Cell Sci 132, (2019).

15. V. Basrur et al., Proteomic analysis of early melanosomes: identification of novel melanosomal proteins. J Proteome Res 2, 69–79 (2003).

16. J. J. Lemasters et al., Compartmentation of Mitochondrial and Oxidative Metabolism in Growing Hair Follicles: A Ring of Fire. The Journal of investigative dermatology 137, 1434–1444 (2017).

17. A. K. McBride, W. F. Bergfeld, Mosaic hair color changes in alopecia areata. Cleve Clin J Med 57, 354–356 (1990).

18. T. Komagamine, K. Suzuki, K. Hirata, Darkening of white hair following levodopa therapy in a patient with Parkinson’s disease. Mov Disord 28, 1643 (2013).

19. N. J. Reynolds, J. Crossley, I. Ferguson, R. D. Peachey, Darkening of white hair in Parkinson’s disease. Clin Exp Dermatol 14, 317–318 (1989).

20. F. Ricci, C. De Simone, L. Del Regno, K. Peris, Drug-induced hair colour changes. Eur J Dermatol 26, 531–536 (2016).

21. B. F. Sieve, Clinical Achromotrichia. Science 94, 257–258 (1941).

22. K. H. Yoon, D. Kim, S. Sohn, W. S. Lee, Segmented heterochromia in scalp hair. J Am Acad Dermatol 49, 1148–1150 (2003).

23. E. Kobayashi et al., Reversible hair depigmentation in a Japanese female treated with pazopanib. J Dermatol 41, 1021–1022 (2014).

24. A. Kavak, Y. Akcan, U. Korkmaz, Hair repigmentation in a hepatitis C patient treated with interferon and ribavirin. Dermatology 211, 171–172 (2005).

25. R. Paus, E. A. Langan, S. Vidali, Y. Ramot, B. Andersen, Neuroendocrinology of the hair follicle: principles and clinical perspectives. Trends Mol Med 20, 559–570 (2014).

26. A. A. Navarini, R. M. Trueb, Reversal of canities. Arch Dermatol 146, 103–104 (2010).

27. S. Comaish, White scalp hairs turning black--an unusual reversal of the ageing process. Br J Dermatol 86, 513–514 (1972).

28. D. J. Tobin, J. A. Cargnello, Partial reversal of canities in a 22-year-old normal Chinese male. Arch Dermatol 129, 789–791 (1993).

29. D. J. Tobin, R. Paus, Graying: gerontobiology of the hair follicle pigmentary unit. Exp Gerontol 36, 29–54 (2001).

30. J. N. O’Sullivan, C.; Picard, M.; Chéret, J.; Bedogni, B.; Tobin, DJ.; Paus, R., The biology of human hair greying. Biol Rev (2020).

31. E. Puterman et al., Lifespan adversity and later adulthood telomere length in the nationally representative US Health and Retirement Study. Proc Natl Acad Sci U S A 113, E6335–E6342 (2016).

32. E. S. Epel et al., Accelerated telomere shortening in response to life stress. Proc Natl Acad Sci U S A 101, 17312–17315 (2004).

33. B. Zhang et al., Hyperactivation of sympathetic nerves drives depletion of melanocyte stem cells. Nature 577, 676–681 (2020).

34. M. A. LeBeau, M. A. Montgomery, J. D. Brewer, The role of variations in growth rate and sample collection on interpreting results of segmental analyses of hair. Forensic Sci Int 210, 110–116 (2011).

35. A. E. Douglass, in Climatic Cycles and Tree Growth. (Carnegie Institute of Washington, Washington, DC, 1928), vol. vol. II..

36. R. Paus, G. Cotsarelis, The biology of hair follicles. N Engl J Med 341, 491–497 (1999).

37. A. Slominski et al., Hair follicle pigmentation. The Journal of investigative dermatology 124, 13–21 (2005).

38. S. B. Cho, Z. Zheng, J. Y. Kim, S. H. Oh, Segmented heterochromia in a single scalp hair. Acta Derm Venereol 94, 609–610 (2014).

39. D. J. Tobin, Aging of the hair follicle pigmentation system. Int J Trichology 1, 83–93 (2009).

40. R. N. Franklin et al., Proteomic genotyping: Using mass spectrometry to infer SNP genotypes in pigmented and non-pigmented hair. Forensic Sci Int 310, 110200 (2020).

41. D. Van Neste, Thickness, medullation and growth rate of female scalp hair are subject to significant variation according to pigmentation and scalp location during ageing. European Journal of Dermatology 14, 28–32 (2004).

42. S. S. Adav et al., Studies on the Proteome of Human Hair - Identification of Histones and Deamidated Keratins. Sci Rep 8, 1599 (2018).

43. S. E. Calvo, K. R. Clauser, V. K. Mootha, MitoCarta2.0: an updated inventory of mammalian mitochondrial proteins. Nucleic Acids Res 44, D1251–1257 (2016).

44. B. Singh, T. R. Schoeb, P. Bajpai, A. Slominski, K. K. Singh, Reversing wrinkled skin and hair loss in mice by restoring mitochondrial function. Cell Death Dis 9, 735 (2018).

45. S. Vidali et al., Hypothalamic-pituitary-thyroid axis hormones stimulate mitochondrial function and biogenesis in human hair follicles. The Journal of investigative dermatology 134, 33–42 (2014).

46. J. E. Kloepper et al., Mitochondrial function in murine skin epithelium is crucial for hair follicle morphogenesis and epithelial-mesenchymal interactions. J Invest Dermatol 135, 679–689 (2015).

47. M. Silengo et al., Hair anomalies as a sign of mitochondrial disease. Eur J Pediatr 162, 459–461 (2003).

48. S. K. Singh et al., E-cadherin mediates ultraviolet radiation- and calcium-induced melanin transfer in human skin cells. Exp Dermatol 26, 1125–1133 (2017).

49. J. M. Wood et al., Senile hair graying: H2O2-mediated oxidative stress affects human hair color by blunting methionine sulfoxide repair. FASEB J 23, 2065–2075 (2009).

50. J. A. Hardman et al., The peripheral clock regulates human pigmentation. The Journal of investigative dermatology 135, 1053–1064 (2015).

51. D. J. Tobin, S. Kauser, Hair melanocytes as neuro-endocrine sensors--pigments for our imagination. Mol Cell Endocrinol 243, 1–11 (2005).

52. E. Gaspar et al., Thyrotropin-releasing hormone selectively stimulates human hair follicle pigmentation. The Journal of investigative dermatology 131, 2368–2377 (2011).

53. M. Nahm, A. A. Navarini, E. W. Kelly, Canities subita: a reappraisal of evidence based on 196 case reports published in the medical literature. Int J Trichology 5, 63–68 (2013).

54. L. K. M. Han et al., Accelerating research on biological aging and mental health: Current challenges and future directions. Psychoneuroendocrinology 106, 293–311 (2019).

55. M. Picard et al., A Mitochondrial Health Index Sensitive to Mood and Caregiving Stress. Biol Psychiatry 84, 9–17 (2018).

56. M. Picard, B. S. McEwen, Psychological Stress and Mitochondria: A Systematic Review. Psychosom Med 80, 141–153 (2018).

57. I. R. Schlaepfer, M. Joshi, CPT1A-mediated Fat Oxidation, Mechanisms, and Therapeutic Potential. Endocrinology 161, (2020).

58. C. Bekeova et al., Multiple mitochondrial thioesterases have distinct tissue and substrate specificity and CoA regulation, suggesting unique functional roles. J Biol Chem 294, 19034–19047 (2019).

59. A. Okado-Matsumoto, I. Fridovich, Subcellular distribution of superoxide dismutases (SOD) in rat liver: Cu,Zn-SOD in mitochondria. J Biol Chem 276, 38388–38393 (2001).

60. L. Hoffmann, M. B. Rust, C. Culmsee, Actin(g) on mitochondria - a role for cofilin1 in neuronal cell death pathways. Biol Chem 400, 1089–1097 (2019).

61. K. Rehklau et al., Cofilin1-dependent actin dynamics control DRP1-mediated mitochondrial fission. Cell Death Dis 8, e3063 (2017).

62. H. Nie et al., O-GlcNAcylation of PGK1 coordinates glycolysis and TCA cycle to promote tumor growth. Nat Commun 11, 36 (2020).

63. G. J. Sung et al., Targeting CPT1A enhances metabolic therapy in human melanoma cells with the BRAF V600E mutation. Int J Biochem Cell Biol 81, 76–81 (2016).

64. V. N. Sumantran, P. Mishra, N. Sudhakar, Microarray analysis of differentially expressed genes regulating lipid metabolism during melanoma progression. Indian J Biochem Biophys 52, 125–131 (2015).

65. C. T. Oh et al., Superoxide dismutase 1 inhibits alpha-melanocyte stimulating hormone and ultraviolet B-induced melanogenesis in murine skin. Ann Dermatol 26, 681–687 (2014).

66. C. Bracalente et al., Cofilin-1 levels and intracellular localization are associated with melanoma prognosis in a cohort of patients. Oncotarget 9, 24097–24108 (2018).

67. D. Morvan, J. M. Steyaert, L. Schwartz, M. Israel, A. Demidem, Normal human melanocytes exposed to chronic insulin and glucose supplementation undergo oncogenic changes and methyl group metabolism cellular redistribution. Am J Physiol Endocrinol Metab 302, E1407–1418 (2012).

68. C. D. Wiley, J. Campisi, From Ancient Pathways to Aging Cells-Connecting Metabolism and Cellular Senescence. Cell Metab 23, 1013–1021 (2016).

69. J. Halloy, B. A. Bernard, G. Loussouarn, A. Goldbeter, Modeling the dynamics of human hair cycles by a follicular automaton. Proc Natl Acad Sci U S A 97, 8328–8333 (2000).

70. E. D. Kinzina, D. I. Podolskiy, S. E. Dmitriev, V. N. Gladyshev, Patterns of Aging Biomarkers, Mortality, and Damaging Mutations Illuminate the Beginning of Aging and Causes of Early-Life Mortality. Cell Rep 29, 4276–4284 e4273 (2019).

71. R. Bahar et al., Increased cell-to-cell variation in gene expression in ageing mouse heart. Nature 441, 1011–1014 (2006).

72. M. Enge et al., Single-Cell Analysis of Human Pancreas Reveals Transcriptional Signatures of Aging and Somatic Mutation Patterns. Cell 171, 321–330 e314 (2017).

73. C. P. Martinez-Jimenez et al., Aging increases cell-to-cell transcriptional variability upon immune stimulation. Science 355, 1433–1436 (2017).

74. K. V. Patel et al., Red cell distribution width and mortality in older adults: a meta-analysis. J Gerontol A Biol Sci Med Sci 65, 258–265 (2010).

75. K. S. Stenn, R. Paus, Controls of hair follicle cycling. Physiol Rev 81, 449–494 (2001).

76. C. R. Robbins, The Chemical and Physical Behavior of Human Hair. (Springer, Berlin, 2012).

77. R. S. Balaban, S. Nemoto, T. Finkel, Mitochondria, oxidants, and aging. Cell 120, 483–495 (2005).

78. M. G. Vizioli et al., Mitochondria-to-nucleus retrograde signaling drives formation of cytoplasmic chromatin and inflammation in senescence. Genes Dev 34, 428–445 (2020).

79. B. K. Kennedy et al., Geroscience: linking aging to chronic disease. Cell 159, 709–713 (2014).

80. A. Latorre-Pellicer et al., Mitochondrial and nuclear DNA matching shapes metabolism and healthy ageing. Nature 535, 561–565 (2016).

81. J. Y. Jang, A. Blum, J. Liu, T. Finkel, The role of mitochondria in aging. J Clin Invest 128, 3662–3670 (2018).

82. G. Mancino et al., The Thyroid Hormone Inactivator Enzyme, Type 3 Deiodinase, Is Essential for Coordination of Keratinocyte Growth and Differentiation. Thyroid, (2020).

83. Y. Ramot et al., Spermidine promotes human hair growth and is a novel modulator of human epithelial stem cell functions. PLoS One 6, e22564 (2011).

84. S. A. Villeda et al., Young blood reverses age-related impairments in cognitive function and synaptic plasticity in mice. Nat Med 20, 659–663 (2014).

85. J. Rebo et al., A single heterochronic blood exchange reveals rapid inhibition of multiple tissues by old blood. Nat Commun 7, 13363 (2016).

86. M. Mehdipour et al., Rejuvenation of three germ layers tissues by exchanging old blood plasma with saline-albumin. Aging (Albany NY) 12, 8790–8819 (2020).

87. E. Puterman et al., Aerobic exercise lengthens telomeres and reduces stress in family caregivers: A randomized controlled trial - Curt Richter Award Paper 2018. Psychoneuroendocrinology 98, 245–252 (2018).

88. G. M. Fahy et al., Reversal of epigenetic aging and immunosenescent trends in humans. Aging Cell 18, e13028 (2019).

89. A. Gilhar, T. Pillar, M. David, Aged versus young skin before and after transplantation onto nude mice. Br J Dermatol 124, 168–171 (1991).

90. A. Gilhar, T. Pillar, M. David, S. Eidelman, Melanocytes and Langerhans cells in aged versus young skin before and after transplantation onto nude mice. J Invest Dermatol 96, 210–214 (1991).

91. T. J. LaRocca, A. N. Cavalier, D. Wahl, Repetitive elements as a transcriptomic marker of aging: Evidence in multiple datasets and models. Aging Cell 19, (2020).

92. J. M. van Deursen, The role of senescent cells in ageing. Nature 509, 439–446 (2014).

93. J. M. Lemons et al., Quiescent fibroblasts exhibit high metabolic activity. PLoS Biol 8, e1000514 (2010).

94. D. J. Baker et al., Naturally occurring p16(Ink4a)-positive cells shorten healthy lifespan. Nature 530, 184–189 (2016).

95. S. Omori et al., Generation of a p16 Reporter Mouse and Its Use to Characterize and Target p16(high) Cells In Vivo. Cell Metab, (2020).

96. J. Franzen et al., Senescence-associated DNA methylation is stochastically acquired in subpopulations of mesenchymal stem cells. Aging Cell 16, 183–191 (2017).

97. O. Kalliokoski, F. K. Jellestad, R. Murison, A systematic review of studies utilizing hair glucocorticoids as a measure of stress suggests the marker is more appropriate for quantifying short-term stressors. Sci Rep 9, 11997 (2019).

