## Supplemental Figures 1-12 for "Quantitative Mapping of Human Hair Graying and Reversal in Relation to Life Stress"

#### Supplemental Figure S1

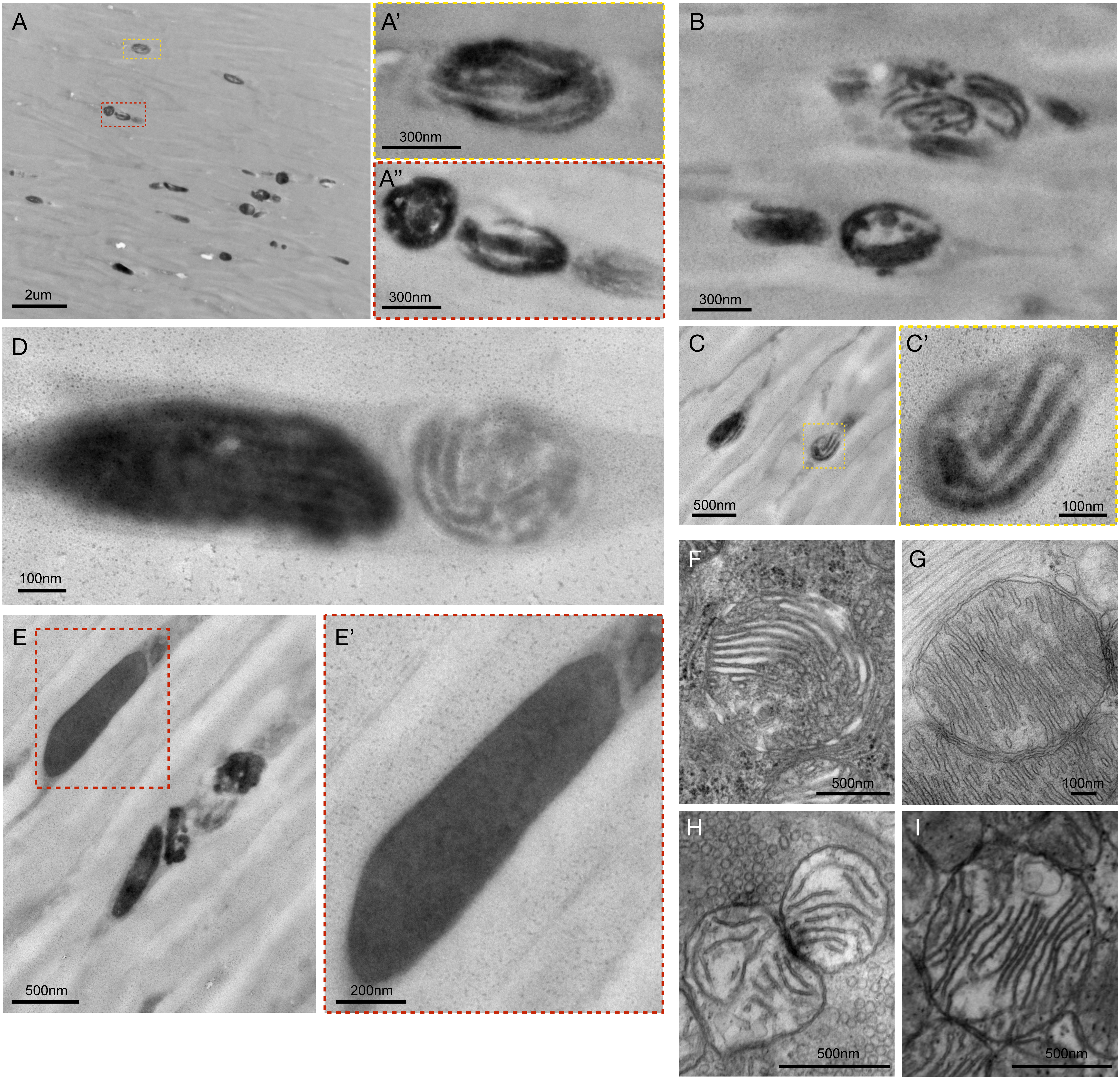


**Figure S1. Ultrastructure of melanosomes and dense granules in human hair shafts.** (**A-D**) Electron micrographs of a dark hair from a 33-year-old Caucasian male with brown hairs. A variety of lamellar melanin granule (melanosomes) are shown. (**E**) Dense amorphous and uniform melanin granules coexist with lamellar melanosomes in the same hair. (**F-I**) Representative images of mitochondria from various tissues showing cristae structure not unlike what is typically ascribed as lamellar melanosomes in the hair. Mitochondria from the mouse adrenal gland (F), heart (G), presynaptic bouton of the neuromuscular junction (H), and skeletal muscle (I) are shown for illustrative purposes. On the basis of available evidence, it cannot be ruled out that some of the pigmented lamellar structures in the HS are remnants of mitochondria and mito-lysosomal hybrids.

### Supplemental Figure S2

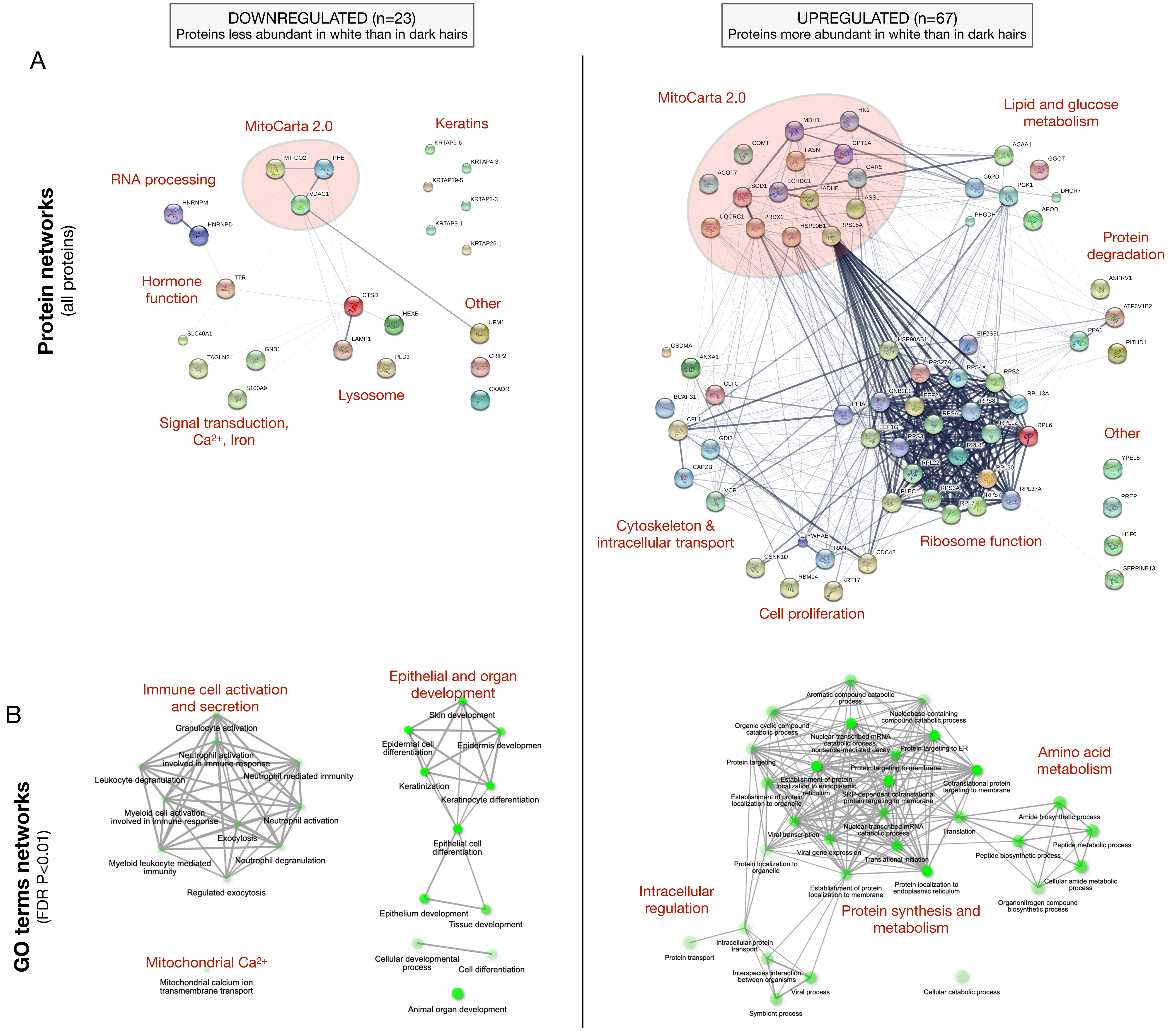


**Figure S2. Functional enrichment analysis of up- and down-regulated proteins in white hair shafts relative to dark from Experiment 1.** (**A**) Protein-protein interaction (PPI) networks from the STRING database for all upregulated (n=67) and down regulated (n=23) proteins. Nodes are proteins, edges reflect evidence of PPI where the thickness reflects the strength of evidence for protein-protein interaction (PPI). (**B**) Top gene ontology (GO) terms for the proteins shown in (A), illustrating various cellular processes overrepresented in each gene set. Nodes are GO categories, edges reflect similarity between GO categories (overlap in the genes that compose them). The size & intensity of nodes reflect the significance in the enrichment score, respectively.

#### Supplemental Figure S3

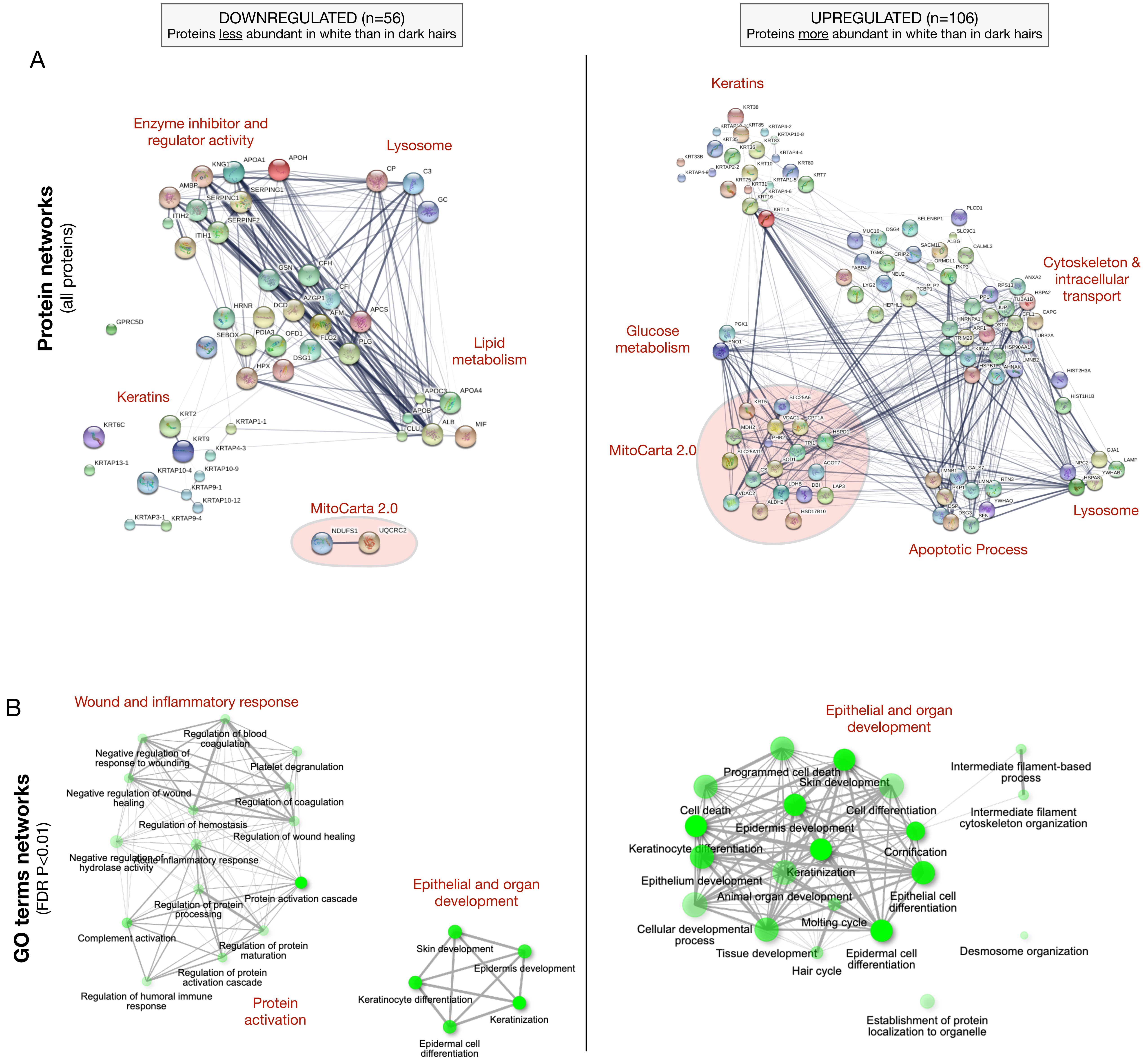


**Figure S3. Functional enrichment analysis of up- and down-regulated proteins in white hair shafts relative to dark in experiment 2.** (**A**) PPI networks from the STRING database for all upregulated (n=106) and down regulated (n=56) proteins. Nodes are proteins, edges reflect evidence of PPI with the thickness reflecting to the strength of evidence. (**B**) Top gene ontology (GO) terms for the proteins shown in (A), illustrating various cellular processes overrepresented in each gene set. Nodes are GO categories, edges reflect similarity between GO categories based on overlap in genes, and size & intensity of nodes reflect the significance in the enrichment scores, respectively.

### Supplemental Figure S4

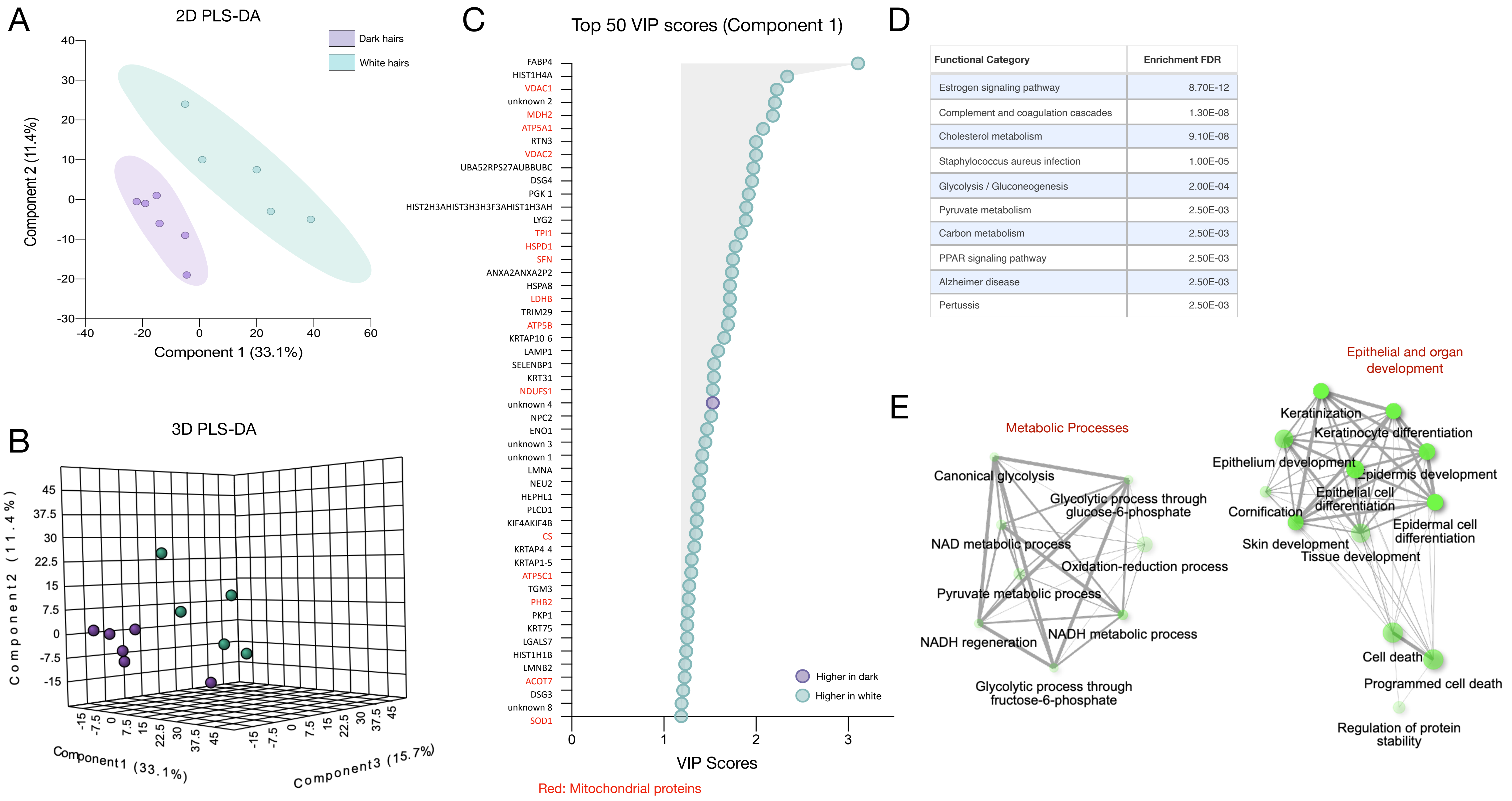


| Functional Category | Enrichment FDR |
|---|---|
| Estrogen signaling pathway | 8.70E-12 |
| Complement and coagulation cascades | 1.30E-08 |
| Cholesterol metabolism | 9.10E-08 |
| Staphylococcus aureus infection | 1.00E-05 |
| Glycolysis / Gluconeogenesis | 2.00E-04 |
| Pyruvate metabolism | 2.50E-03 |
| Carbon metabolism | 2.50E-03 |
| PPAR signaling pathway | 2.50E-03 |
| Alzheimer disease | 2.50E-03 |
| Pertussis | 2.50E-03 |

**Figure S4. Partial least square discriminant analysis (PLS-DA) of dark and white hairs from experiment 2.** (**A**) 2D and (**B**) 3D score plot of dark and white HS generated from protein abundance (n=192) quantified by LC-MS/MS. (**C**) The top 50 protein that maximally contribute to group separation ranked by their variable importance in projection (VIP) scores on the model's first component, colored by up or down-regulation status in white HS relative to dark. Note that 49 of 50 proteins (98%) are upregulated (more abundant) in white hairs relative to dark. (**D**) KEGG functional annotation categories with enrichment p values. (**E**) GO term networks indicating coherent groups of categories related to metabolic processes involving NAD/NADH metabolic processes and substrate oxidation, as well as epithelial and organ development categories, including cell death. Nodes are GO categories, edges reflect similarity between GO categories based on overlap in genes, and size & intensity of nodes reflect the significance in the enrichment scores, respectively.

#### Supplemental Figure S5

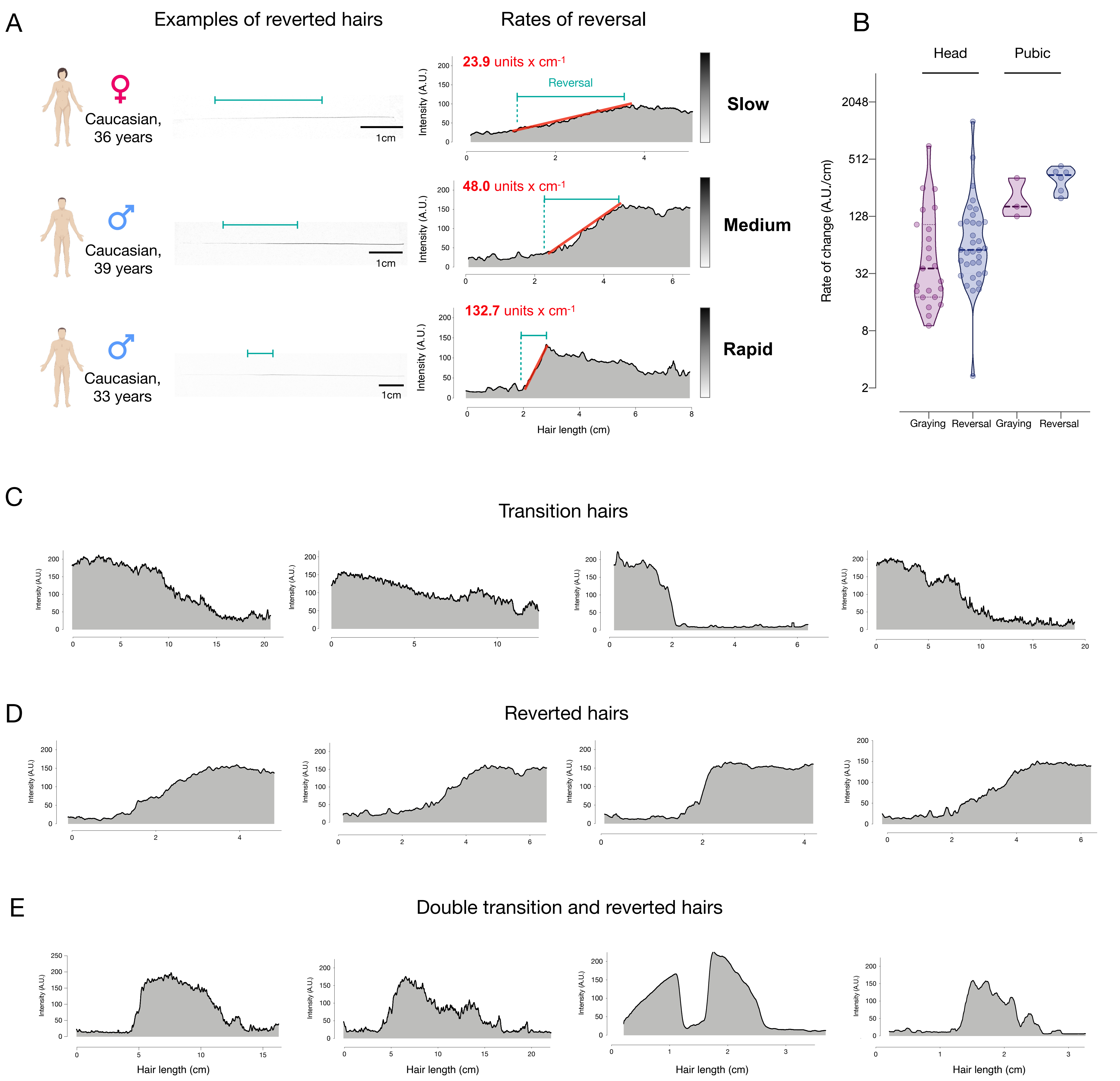


**Figure S5. Hair pigmentation patterns across a range of individuals.** (**A**) Examples of graying reversal in hairs with various rates of re-pigmentation in three adult individuals, including slow, medium, and rapid events of repigmentation. The left side shows actual photographs of hairs and the length over which each hair shaft regains pigmentation. On the right are the corresponding hair pigmentation patterns (HPPs) for each hair and the computed slope reflecting the rate of graying reversal. (**B**) Quantification of the rate of change in pigmentation per day in graying and reverted hairs (n = 39) and pubic graying and reverted hairs (n = 9), reported on a $\log_2$ scale. (**C**) Examples of hairs undergoing the transition from dark to gray from a 40-year-old female, 35-year-old female, and 69-year-old male. (**D**) Additional examples of hairs undergoing graying reversal from a 39-year-old male, 36-year-old female, and a 33-year-old male. (**E**) Additional examples of hairs with double transitions and reversions from a 35-year-old female and a 36-year-old female.

#### Supplemental Figure S6

### Stress Graph

##### Instructions

*** Please Read Full Instructions on Prior Page Before Completing the Graph.**

1. Mark the **most stressful period** at **10** on the graph.
2. Mark the **least stressful period** at **0** on the graph.
3. Indicate **2-8 stressful periods** by marking them at a point on the graph with a number. Shortly describe it in the associated box on the bottom of the page.
4. Draw a line for the whole year, showing the variation in your experience of **stress** over time.

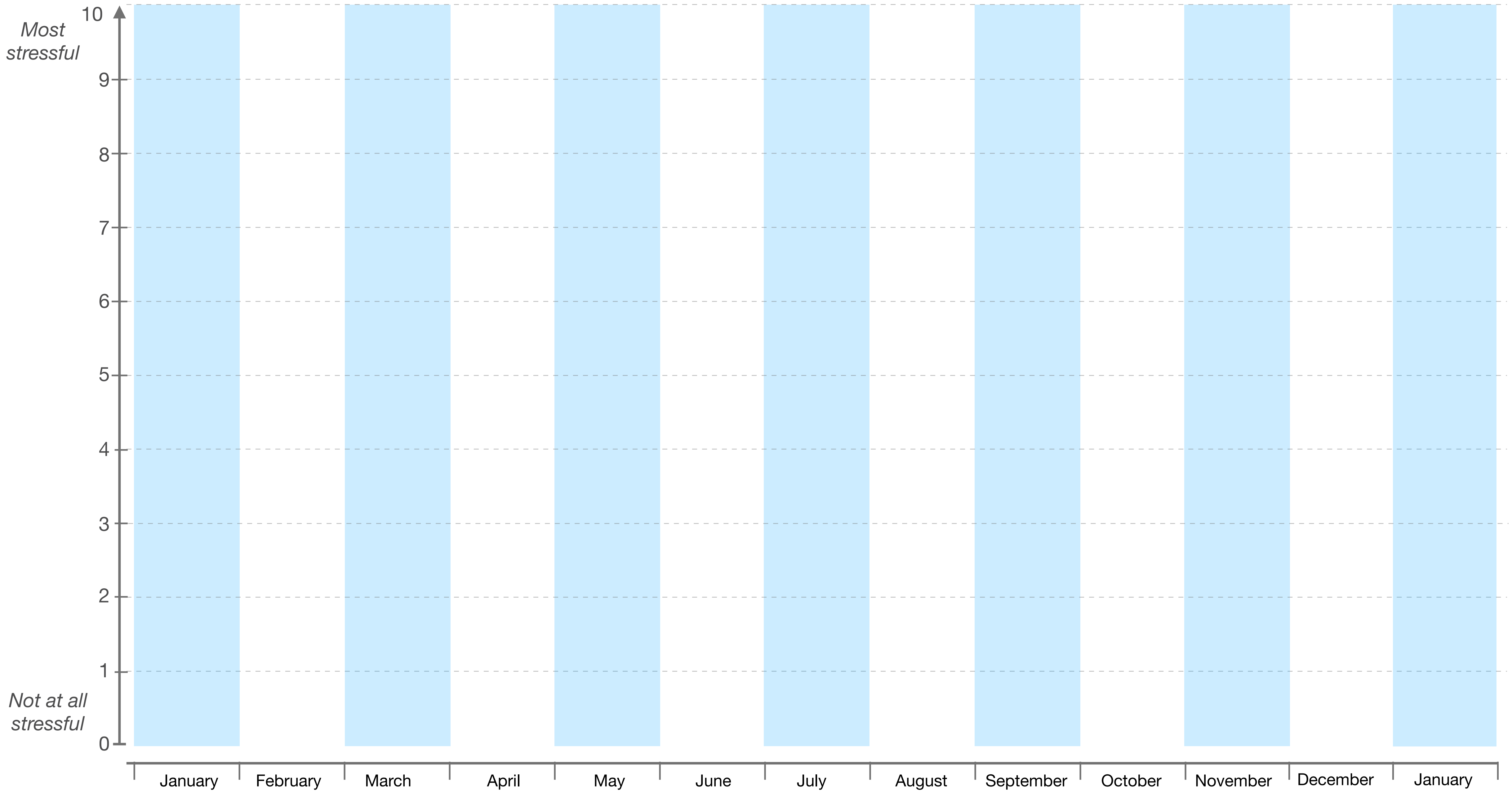


**1. Most Stressful**

**2. Least Stressful**

**3.**

**4.**

**5.**

**6.**

**7.**

**8.**

**Figure S6. Retrospective stress assessment instrument.** Participants are asked to note the most and least stressful events over the past year (Boxes 1 and 2) and mark them with a corresponding point on the scheme above. Participants then indicate 2-6 noteworthy life events or periods over the past year and assign them scores ranging from most stressful to least stressful (10 and 0, respectively) and mark them on the timeline. Participants are asked to briefly name/describe each event, to help with recall. Participants then connect these events with a line. The stress graph is subsequently digitized and used in correlation analyses with hair pigmentation patterns (HPPs) as in Figures 3C and 3D.

### Supplemental Figure S7

A

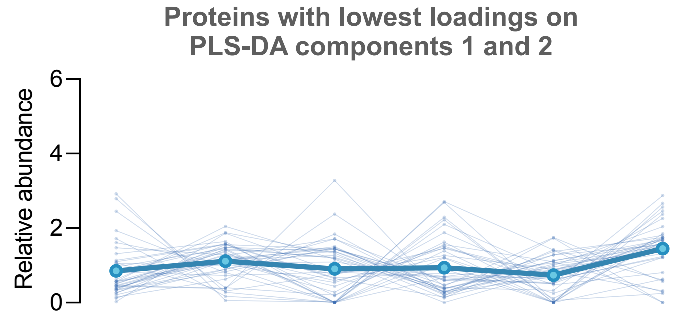


No loading: BACKGROUND

| Functional Category | Enrichment FDR |
|---|---|
| Cholesterol metabolism | 0.000009 |
| Vitamin digestion and absorption | 0.00003 |
| Fat digestion and absorption | 0.0001 |
| Estrogen signaling pathway | 0.0001 |
| Staphylococcus aureus infection | 0.0079 |
| PPAR signaling pathway | 0.0083 |
| Complement and coagulation cascades | 0.0083 |
| Spliceosome | 0.020 |
| Protein processing in endoplasmic reticulum | 0.026 |

B

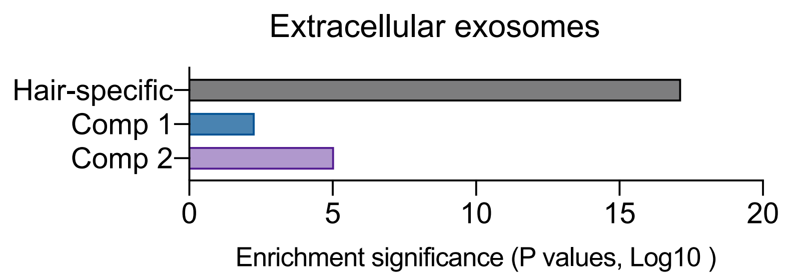


C

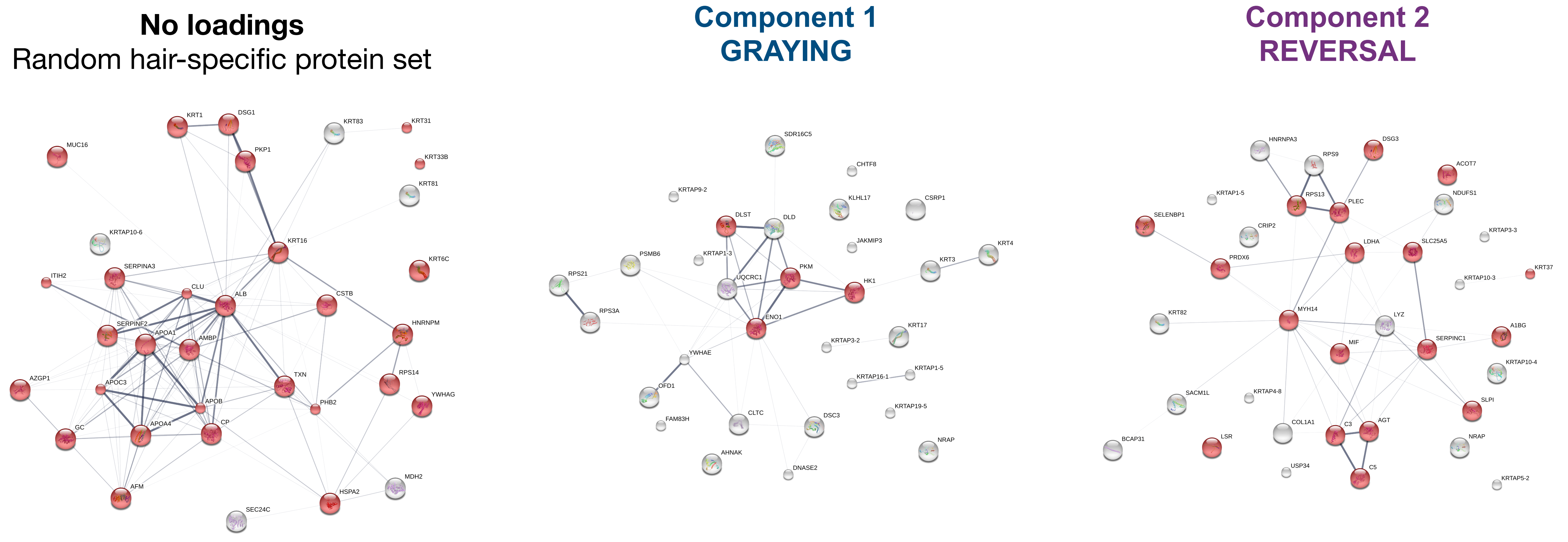


Nodes related to **extracellular exosomes** in red (Top cellular component)

Nodes related to **NADH metabolism** in red (Top molecular function)

Nodes related to **extracellular exosomes** in red (Top cellular component)

**Figure S7. Single-hair multi-segment analysis. (A)** Abundance trajectories of the 20 proteins with lowest loadings on the PLS-DA model components 1 and 2, reflecting background protein composition not related to the graying process. The top KEGG categories enriched among these proteins is listed in the table. (**B**) Enrichment significance for the GO term “extracellular exosomes” among proteins with highest loadings on components 1 and 2 compared with background proteins (lowest loadings) showing a marked decrease in exosome-related signature with graying (component 1), which is more modestly attenuated among component 2 proteins. (**C**) Protein-protein interaction networks (STRING) for background proteins with no loadings, component 1 proteins, and component 2 proteins, highlighting nodes relating to top GO cellular component or molecular function. Note the high number of proteins related to extracellular exosomes in background hair proteins, compared to proteins associated with hair graying. The molecular signature associated with component 2 (reversal) appears to show a re-enrichment for exosome and more closely resembles the non-specific proteins.

### Supplemental Figure S8

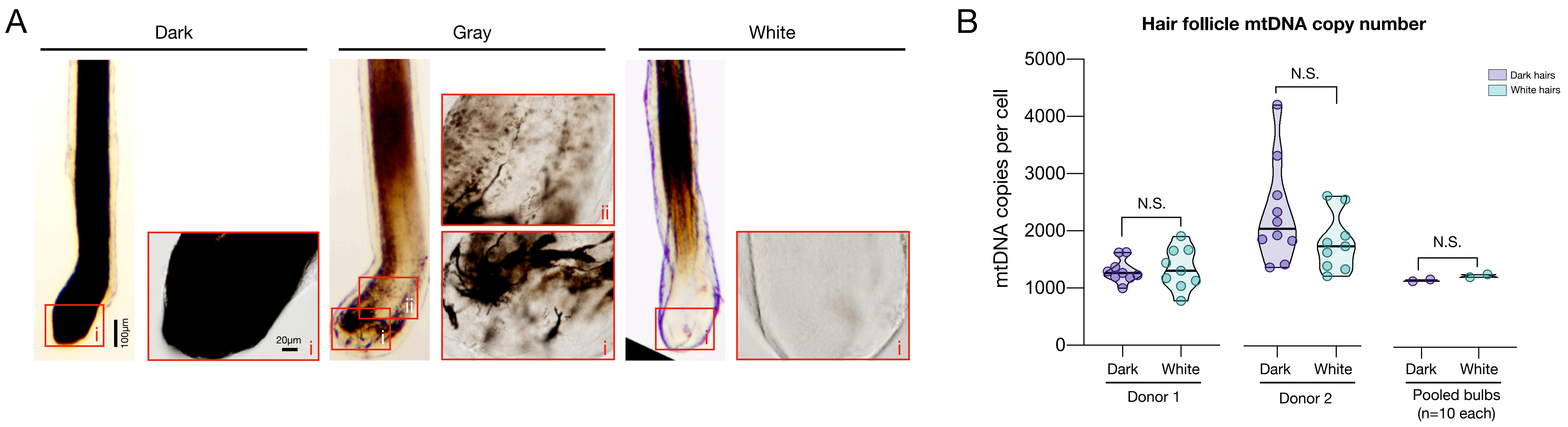


**Figure S8. Mitochondrial DNA copy number in dark and white hair follicles.** (**A**) Typical examples of pigmented (dark), gray, and depigmented (white) plucked hair follicles, which include vital tissue from the precortical hair matrix and at least remnants of the hair follicle pigmentary unit (HFPU) imaged by light microscopy on freshly plucked HFs. Note the absence of pigmented melanocytes in the white HF. Boxed regions show magnified versions for each hair type. (**B**) Mitochondrial and nuclear DNA abundance were quantified from the hair follicles of plucked dark and white hairs, like in (A), from the same donors as the electron microscopy in Figure 1 were (male African-American, male caucasian). n=10 hair follicles per condition. To rule out potential differences related to DNA abundance,10 follicles from each condition (donor and color) were pooled and included in a single PCR run (*right*). There is no evidence of mtDNA copy number (mtDNAcn) alterations in white vs dark hair follicles, similar to shafts shown in Figure 4D. Two-way ANOVA, Tukey's multiple comparison test. N.S.: non-significant.

### Supplemental Figure S9

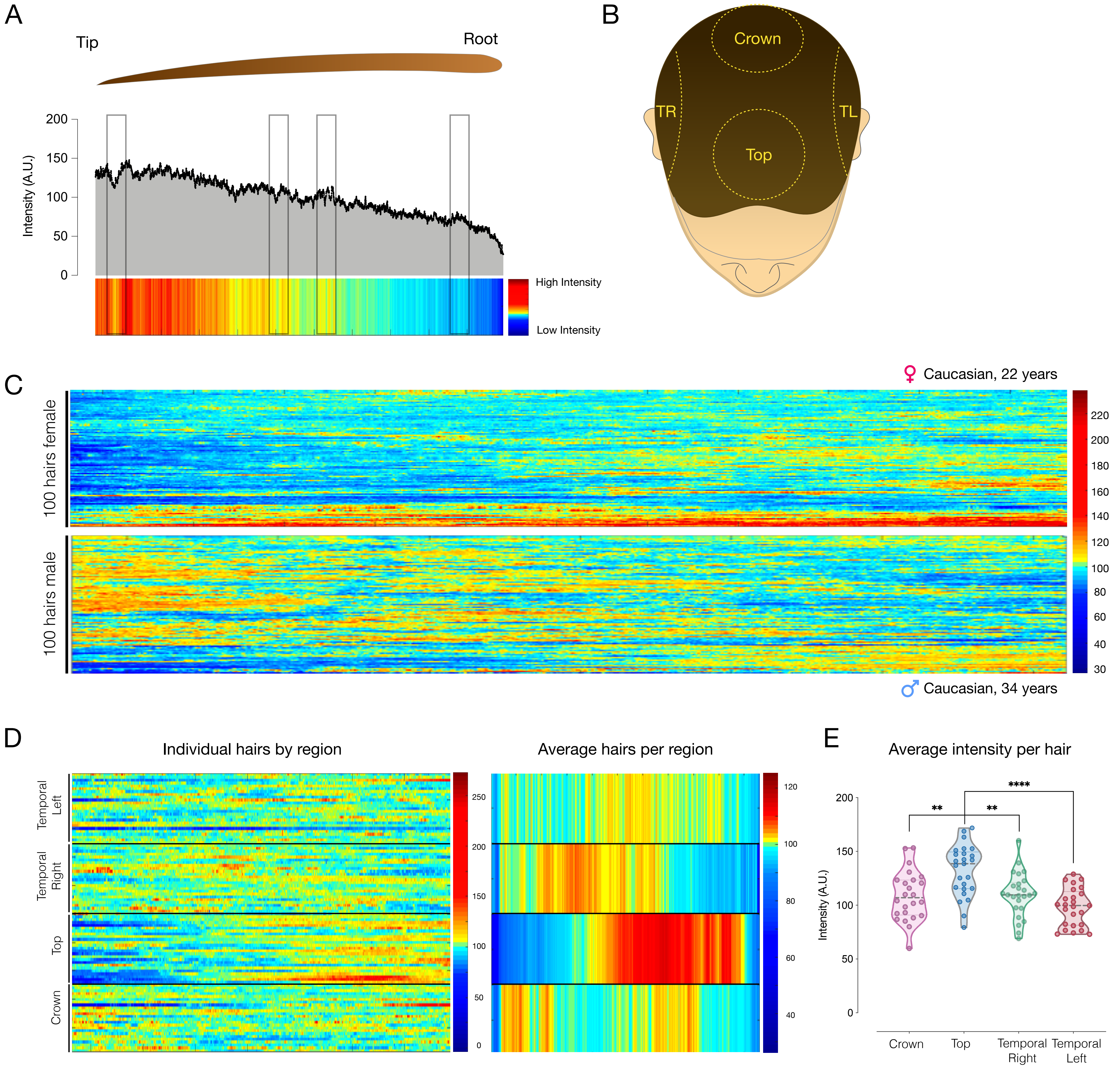


**Figure S9. High-resolution analysis of hair pigmentation patterns.** (**A**) Example of HPP visualized as a heatmap, where dark red represents higher intensity pigmentation values (255 units, black), and dark blue represented lower intensity pigmentation values (20 units, white). The X axis represents the distance from the hair bulb, in pixels, with a resolution of 1,252 pixels/cm. The boxed regions draw attention to some areas along the hair where there are minor changes in HPP, which are captured in the change in color on the heatmap. (**B**) Diagram representing the four regions of the scalp from which hairs were systematically collected; temporal left (TL) and right (TR), top, and crown. (**C**) Hair pigmentation analysis for 100 hairs from an adult female individual (*top*), and adult male individual (*bottom*) from the four head regions shown in B. Hairs are ordered by hierarchical clustering based on Euclidian distance, measuring the similarity of normalized hair pigmentation patterns. Each hair is intensity normalized to reflect deviation from the mean of each hair. (**D**) Same hairs as in (C, Male) but arranged by head regions showing HPP of individual hairs (*left*) or averages for each head region (*right*). (**E**) Average hair darkness (pigmentation intensity) arranged by head regions. Each datapoint is the average of a single hair. **P<0.01, ****P<0.0001, one-way ANOVA, Tukey's multiple comparison test.

### Supplemental Figure S10

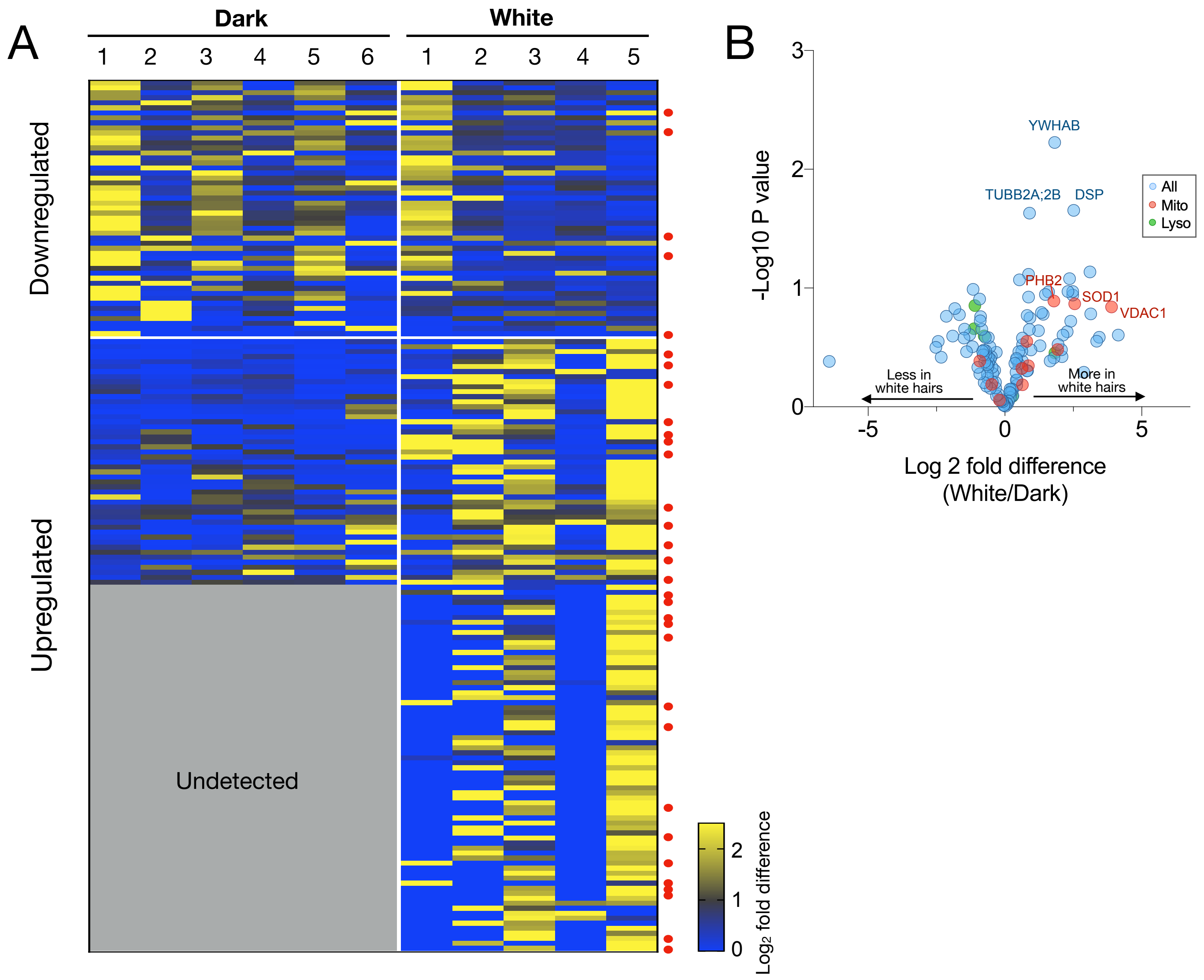


C

##### Upregulated

| Functional Category | Enrichment FDR |
|---|---|
| Estrogen signaling pathway | 4.59E-12 |
| Arrhythmogenic right ventricular cardiomyopathy (ARVC) | 0.00016367 |

##### Downregulated

| Functional Category | Enrichment FDR |
|---|---|
| Keratinization | 4.73E-12 |
| Epidermis development | 3.00E-11 |
| Keratinocyte differentiation | 4.19E-11 |
| Skin development | 6.27E-11 |
| Epidermal cell differentiation | 2.16E-10 |
| Platelet degranulation | 9.97E-10 |
| Regulated exocytosis | 1.67E-09 |
| Exocytosis | 9.62E-09 |
| Chylomicron remodeling | 1.69E-07 |
| Negative regulation of peptidase activity | 2.20E-07 |

**Figure S10. Sensitivity analysis of results from LC-MS:MS hair shaft proteomics experiment 2.** In this analysis, only proteins detected in at least 2/6 dark and 2/5 white hairs were selected. (**A**) Heatmap of down- and up-regulated proteins across all dark and white hairs. Red dots to the left of heatmap indicate mitochondrial proteins, and (**B**) volcano plot of the hair proteome comparing white and dark hairs. (**C**) KEGG pathways functional enrichment analyses for upregulated and downregulated proteins.

### Supplemental Figure S11

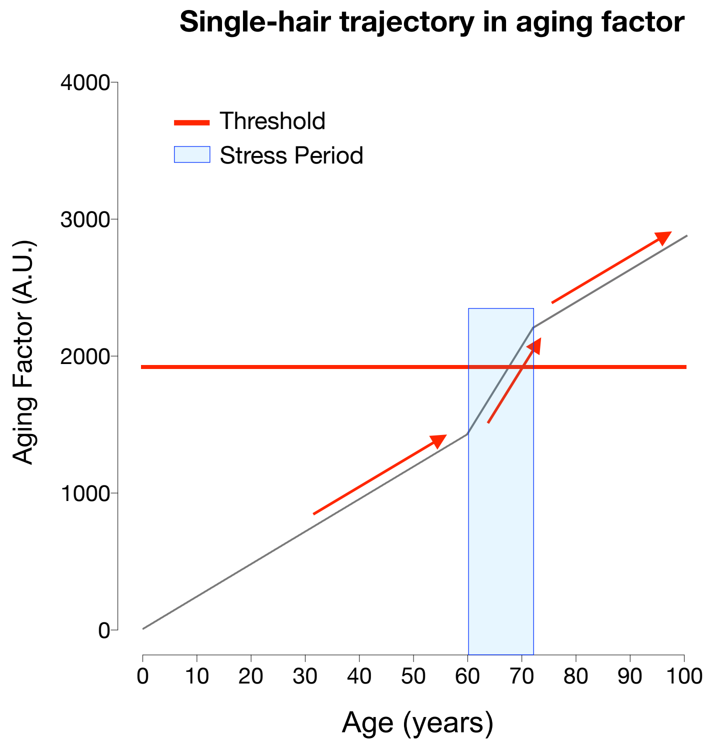


**Figure S11. Alternative modeling of HPP graying transitions in response to stress.** Alternative model examining the influence of stress on the age-related increase in the aging factor. Compared to the successful model where stress causes a stepwise increase in the aging factor, in this model stress increases the slope in the aging factor during a stressful period, and removal of the stressor restores the initial slope of aging factor increase over time. Note that this “changing slope” model fails to account for the reversal of graying upon alleviation of stress, even if the stressor and its removal occur close to the threshold.

#### Supplemental Figure S12

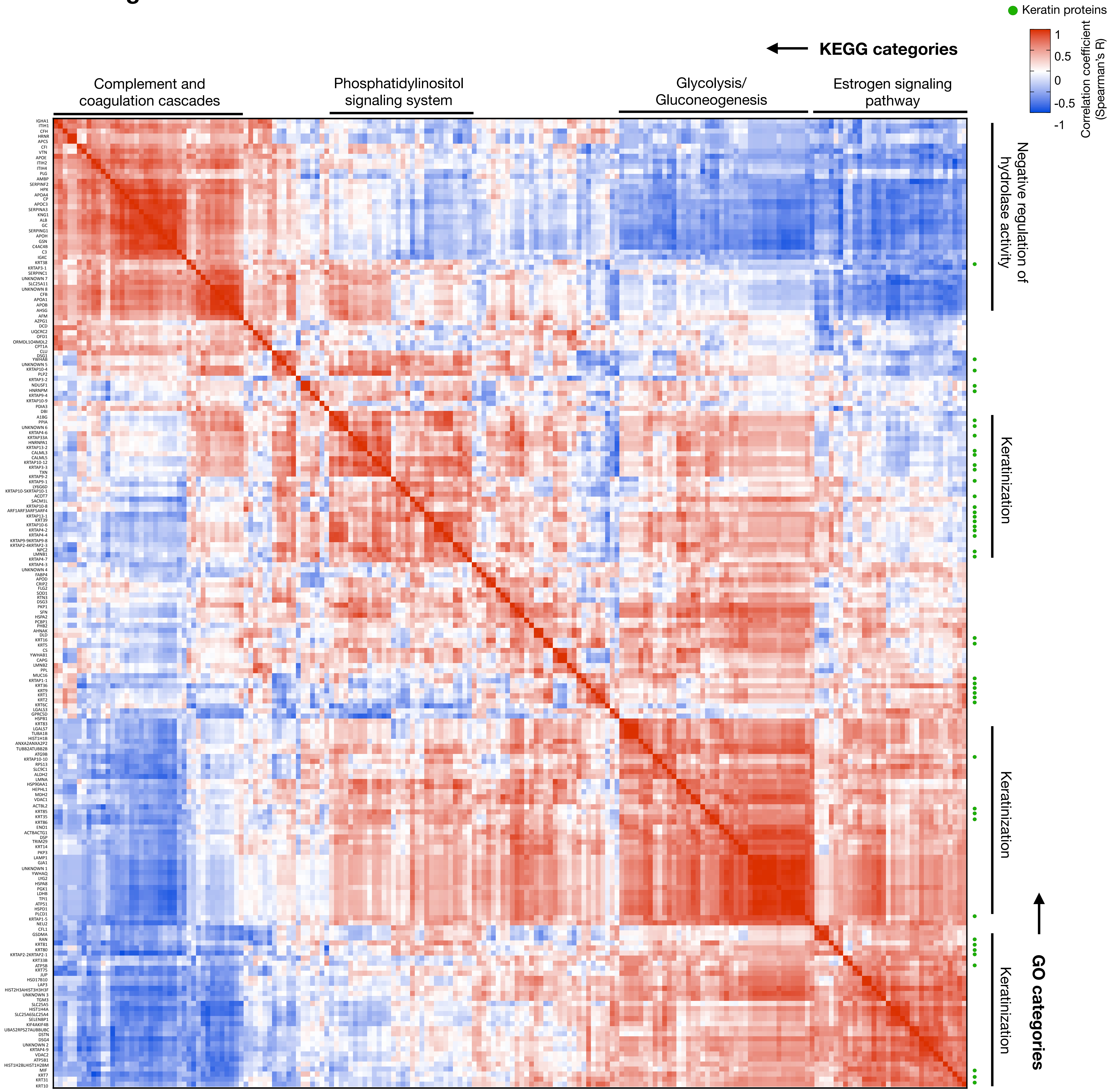


**Figure S12. Correlation matrix reflection co-regulation of the human hair proteome (n=192).** Higher resolution version of Figure 4F. Clusters are labeled by top KEGG (*top*) and GO biological function (*right*) annotations. Correlation strengths and direction are color coded as indicated, and keratin proteins are marked with a green marker. Data from LC-MS:MS experiment 2, n=11 HS segments from 6 individuals.
